# Pore engineering as a general strategy to improve protein-based enzyme nanoreactor performance

**DOI:** 10.1101/2024.05.02.592161

**Authors:** Seokmu Kwon, Michael P. Andreas, Tobias W. Giessen

## Abstract

Enzyme nanoreactors are nanoscale compartments consisting of encapsulated enzymes and a selectively permeable barrier. Sequestration and co-localization of enzymes can increase catalytic activity, stability, and longevity, highly desirable features for many biotechnological and biomedical applications of enzyme catalysts. One promising strategy to construct enzyme nanoreactors is to repurpose protein nanocages found in nature. However, protein-based enzyme nanoreactors often exhibit decreased catalytic activity, partially caused by a mismatch of protein shell selectivity and the substrate requirements of encapsulated enzymes. No broadly applicable and modular protein-based nanoreactor platform is currently available. Here, we introduce a pore-engineered universal enzyme nanoreactor platform based on encapsulins – microbial self-assembling protein nanocompartments with programmable and selective enzyme packaging capabilities. We structurally characterize our protein shell designs via cryo-electron microscopy and highlight their polymorphic nature. Through fluorescence polarization assays, we show their improved molecular flux behavior and highlight their expanded substrate range via a number of proof-of-concept enzyme nanoreactor designs. This work lays the foundation for utilizing our encapsulin-based nanoreactor platform for future biotechnological and biomedical applications.

## Introduction

Protein-based enzyme nanoreactors hold tremendous promise for various applications^1^. This includes both *in vivo* applications related to metabolic engineering^2^ and bioremediation^3^, *in vitro* applications as chemoenzymatic production systems^4^, and applications as delivery devices for biomedically relevant enzymes^5^. Encapsulating enzymes inside nanoreactors can result in a number of distinct advantages, including the controllable co-localization of enzymes^6^, the selective enrichment of substrates^7^, intermediate sequestration and channeling^8^, prevention of unwanted side reactions and toxicity^9^, increased enzyme stability and protease resistance^10–13^, and the ability to sequester and store molecules of interest in a soluble and non-toxic form^14^. Depending on the system, nanoreactors are often assembled *in vitro* using relatively harsh disassembly, mixing, and reassembly protocols^15–18^. All potential nanoreactor applications rely on enzyme encapsulation not reducing enzyme activity below useful levels. So far, this has proved to be challenging to achieve in many nanoreactor systems^4^. More often than not, encapsulated enzymes show substantially decreased activity which we hypothesize is often caused by a mismatch between shell selectivity and the substrate and cofactor requirements of sequestered enzymes.

While various lipid-^19^, polymer-^20^, DNA-^21^, or inorganics^22^-based enzyme nanoreactor designs have been reported in the literature, the necessity for labor intensive *in vitro* manipulations during their construction may pose substantial scale-up and in the end usability challenges. Protein-based nanoreactor designs on the other hand have the potential to be assembled in a single step *in vivo*. In addition, their inherent genetic engineerability may allow for fast design-build-test cycles to optimize designs. So far, a number of different protein-based compartmentalization systems have been used for enzyme nanoreactor applications. These include ferritins^23^, viral capsids^24^, bacterial microcompartments (BMCs)^25^, lumazine synthase^26^, *de novo* designed protein shells^27^, and encapsulins^28^. The P22 viral capsid in particular has been used to create nanoreactors for alcohol oxidation^29^, decarboxylation^30^, hydrogen production^31^, and the degradation of endocrine disruptors^32^. Some of these nanoreactors showed comparable activity with free enzymes, but often exhibited decreased performance. One potential downside of P22 are its large pores of 3 to 10 nm in size depending on the P22 assembly state^33^. This makes the P22 shell quite porous, acting more like an enzyme scaffold than a proper encapsulation system resulting in suboptimal cargo sequestration and protection. Lumazine synthase has also been engineered for nanoreactor applications^26^. This includes a proteasome-like system based on the encapsulation of a protease inside the lumazine synthase shell which exhibited charge-selective peptide degradation behavior but substantially decreased enzymatic activity^34^. Further, lumazine synthase has been employed to engineer a minimal carboxysome-like protein compartment by encapsulating ribulose-1,5-bisphosphate carboxylase/oxygenase (RuBisCO) and carbonic anhydrase (CA) yielding a carbon fixing nanoreactor^35^. It was found that co-encapsulation did not result in improved enzymatic activity. BMCs have been engineered for ethanol^36^ and hydrogen^37^ production, in both cases enzymatic activity decreased upon encapsulation. Due to their robust plug-and-play cargo loading mechanism, encapsulins have also been engineered for nanoreactor applications including for phenanthridine^11^ and 5-methyltetrahydrofolate^38^ production, luciferase encapsulation^12^, and alcohol and oligosaccharides oxidation^13^. However, consistently, as the size of substrates increased, the catalytic activity of encapsulated enzymes significantly decreased.

A number of strategies have been employed to investigate how the properties of pores in protein-based compartments influence molecular flux across protein shells^7,33,37,39–41^. The influence of pore size on shell porosity has been examined for some model systems including the P22 viral capsid^33^, BMCs^7,37^, and encapsulins^38,39^. Consistently across these studies, it has been shown that pore size establishes a size threshold for molecular diffusion where molecules larger than the pore experience significantly hampered molecular flux while molecules smaller than the pore can nearly freely pass through the protein shell. It has also been shown that the charge of pores can influence diffusion across protein shells where molecular flux for molecules or ions with the same charge as the pore is generally hindered due to electrostatic repulsion^33,34,37,39,40^. Measuring molecular flux or protein shell porosity is a non-trivial experimental challenge. Some recently employed approaches include computational flux simulations^40^, lanthanide ion diffusion assays^39^, and the use of NADH-dependent enzymes in combination with differently sized dendrimer-modified NADH derivatives^33^. While innovative and useful, these experimental strategies do not represent ideal model systems for the majority of small molecule enzyme substrates.

Here, we report a modular nanoreactor design based on a pore-engineered encapsulin nanocompartment (Mx_pmut) with optimized shell porosity and molecular flux behavior. We structurally characterize our new shell design using cryo-electron microscopy (cryo-EM) and highlight its polymorphic nature and high cargo loading capacity. Using fluorescence polarization assays, we characterize molecular flux across the Mx_pmut shell. We construct a number of enzyme-loaded proof-of-concept nanoreactors, including multi-enzyme systems, with overall improved catalytic performance. Our work lays the foundation for a universal enzyme nanoreactor platform with potential applications in biomedicine, biocatalysis, and bionanotechnology.

## Results

### Rational pore design as a general strategy for improving enzyme nanoreactor performance

To address the often-observed issue of decreased enzyme activity upon encapsulation inside a protein shell, we will pursue a pore engineering strategy focused on increasing shell porosity. By increasing pore size, we aim to optimize molecular flux of a broad range of enzyme substrates and cofactors across the protein shell. At the same time, our pore design will take into account that very large pores (>3 nm) can lead to the loss of many advantages of enzyme sequestration. This includes the loss of increased local substrate and intermediate concentrations, the loss of protection from proteases, and the loss of the exclusion of unwanted proteinaceous components from the interior of the protein shell. Our nanoreactor design strategy will focus on encapsulins, self-assembling icosahedral protein nanocompartments that range in size from 24 to 42 nm.^42^ They exhibit different triangulation numbers (T=1: 60 subunits, T=3: 180 subunits, or T=4: 240 subunits) and form via self-assembly of a single type of shell protein subunit.^43–46^ Encapsulins generally possess primary pores at the center of all shell pentamers and hexamers ranging in size from 1 to 20 Å.^42^ However, larger pore sizes (>8 Å) have only been observed in select T=1 systems that exhibit dynamic pentameric pores with distinct closed and open states.^47^ In general, T=1 shells possess 12 pores, T=3 shells 32 pores, and T=4 shells 42 pores. The eponymous feature of encapsulins is their ability to selectively encapsulate dedicated cargo proteins during shell self-assembly.^43,48,49^ Encapsulation is facilitated by an efficient cargo loading mechanism based on a C– or N-terminal targeting peptide (TPs) or domain present in all native cargo proteins. Encapsulins possess a number of promising properties for nanoreactor engineering. Firstly, encapsulin systems natively possess a modular and highly effective *in vivo* encapsulation mechanism with which to target essentially any protein of interest to the shell interior. Secondly, encapsulin shells have proved to be highly engineerable and robust, opening up the possibility of advanced nanoreactor applications.^50–52^ Thirdly, encapsulin shell porosity is controlled by engineerable pores.^39,47^ We hypothesize that optimizing encapsulin pores for small molecule flux while not creating protein shells that are too porous will result in a broadly useful nanoreactor platform.

While a small number of pore-engineered encapsulin-based nanoreactor designs – all using small T=1 shells – have been reported^38–40^, they often suffered from low solubility, flexible and less defined pores impeding molecular flux, and importantly, limited cargo loading capacity. Our nanoreactor design is based on a model T=3 encapsulin system from *Myxococcus xanthus* (Mx_wt) with a shell diameter of 32 nm. Mx_wt has twenty three-fold and twelve five-fold pores located at the center of all hexameric and pentameric shell facets, respectively (Fig. 1a). Based on the closest distance of two atoms across the pores, measured in the Mx_wt atomic shell model, three-fold pores are 3.6 Å wide while five-fold pores exhibit a diameter of 6.4 Å. However, Monte Carlo-based simulations using HOLE^53^ yielded substantially smaller values (3-fold: 0.8 Å, 5-fold: 3.4 Å), as has been previously noted for pore mutants of the *Thermotoga maritima* T=1 encapsulin (Fig. 1b,c). Our pore engineering strategy entails deletion of the eight-residue loop (I_195_YEKTGVL_202_) connecting helices α6 and α7 in the A-domain of Mx_wt which should yield a pore-engineered construct – Mx_pmut – with increased pore size. At the same time, the charged residues E_197_ and K_198_ contained with the α6-α7 loop will not be present in the resulting Mx_pmut system, thus decreasing the potential for unfavorable electrostatic interactions during small molecule transit through Mx_pmut shells (Fig. 1d). This loop was chosen because it forms the central contact point within pentameric and hexameric shell facets, forming the respective pores. In addition to deleting loop residues, we will introduce two glycine residues to allow for the necessary tight turn between helices α6 and α7 without disturbing shell protein folding and assembly. Unlike T=1 encapsulins, which only consist of pentameric facets, T=3 and T=4 encapsulins additionally contain hexametric facets. Based on previous reports, hexameric facets are prone to losing their structural integrity upon deletion of pore forming residues prohibiting the formation of stable hexamers and resulting in all-pentamer T=1 shells with limited cargo loading capacity^54^. Therefore, stabilizing both pore-mutated hexametric and pentameric facets simultaneously is a prerequisite for the formation of large T=3 and T=4 shells. We reasoned that this stabilization could be achieved through co-expression of cargo proteins to act as an internal scaffold expanding shells to larger, defined, and more stable assemblies. As expected, without cargo co-expression, Mx_pmut only formed ca. 20 nm T=1 shells as judged by negative stain transmission electron microscopy (TEM) analysis (Fig. 1e and Supplementary Fig. 1 and 2). In contrast, cargo co-expression resulted in highly cargo-loaded – 97% based on gel densitometry – non-T=1 shells with diameters between 35 and 40 nm (Fig. 1f and Supplementary Fig. 1 and 2). As an initial cargo, we chose the small (19 kDa) and monomeric self-labelling protein SNAP-tag carrying an Mx-targeting peptide (TP) at its C-terminus (PEKRLTVGSLRR), which would also be utilized in our downstream molecular flux experiments.

**Fig 1.**
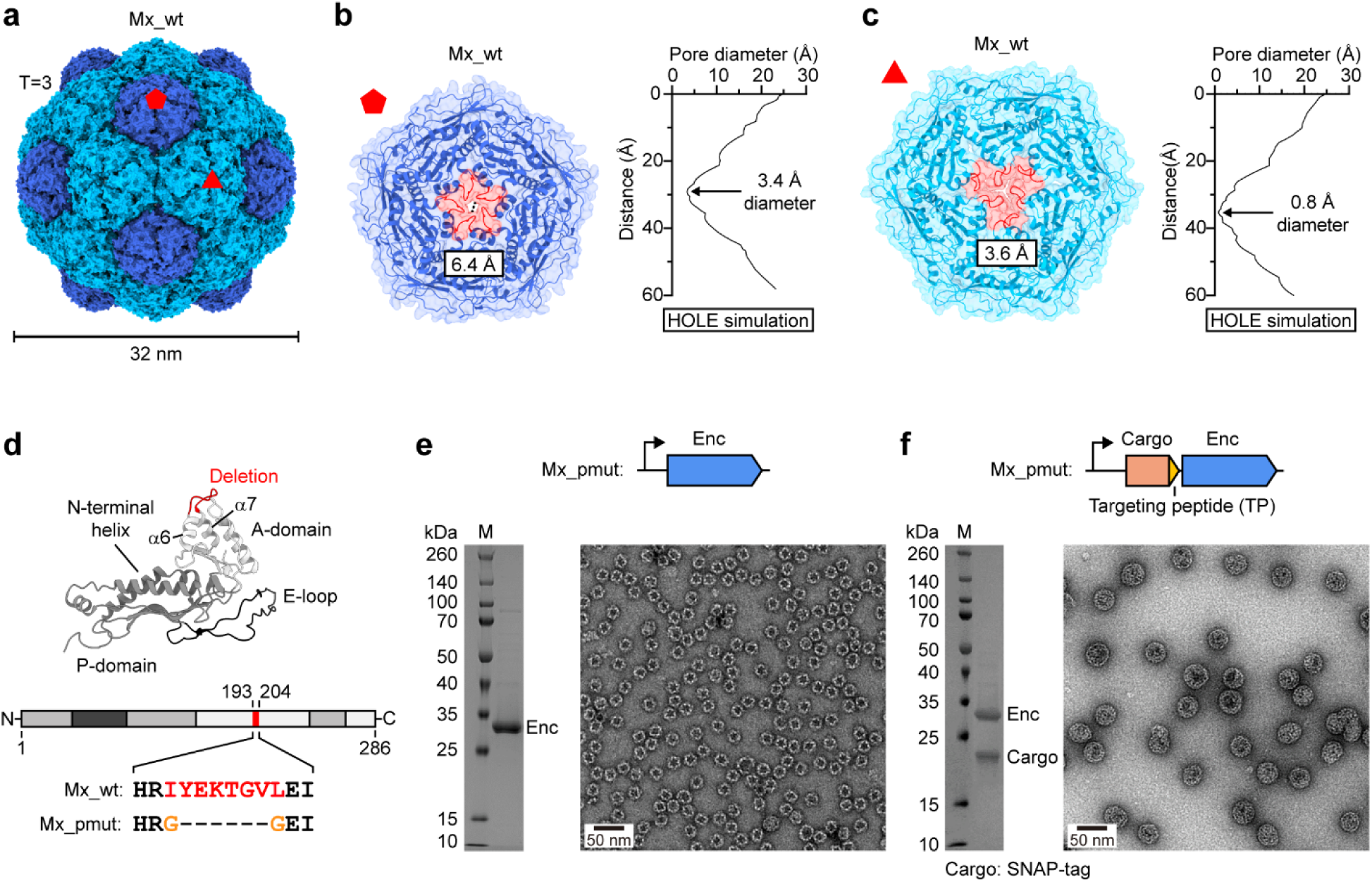
Design and characterization of pore-engineered encapsulin shells. **a**, Native T=3 Mx_wt encapsulin shell. One five– and one three-fold pore are highlighted at the center of a pentameric (dark blue) and a hexameric (cyan) facet, respectively. **b**, Left: Close-up exterior view down the five-fold axis of symmetry onto the five-fold pore of Mx_wt. Pore diameter as measured based on the atomic model is shown. The pore loop to be deleted for our pore-engineered design is highlighted in red. Right: Mx_wt five-fold pore diameter profile as derived from HOLE simulations highlighting the narrowest point of the pore. **c**, Left: Close-up exterior view down the three-fold axis of symmetry onto the three-fold pore of Mx_wt. Pore diameter as measured based on the atomic model is shown. The pore loop to be deleted for our pore-engineered design is highlighted in red. Right: Mx_wt three-fold pore diameter profile as derived from HOLE simulations highlighting the narrowest point of the pore. **d**, Mx_wt protomer with highlighted loop deletion (red) connecting A-domain helices α6 and α7 (top). Pore region sequences of Mx_wt and Mx_pmut showing deleted (red) and added (orange) residues (bottom). **e**, SDS-PAGE analysis and negative-stain TEM micrograph of purified Mx_pmut shells without cargo protein co-expression. Without cargo, Mx_pmut exclusively forms T=1 shells. **f**, SDS-PAGE analysis and negative-stain TEM micrograph of purified Mx_pmut with co-expressed SNAP-tag, carrying a targeting peptide (TP), as a cargo protein. When co-expressed with cargo, Mx_pmut forms larger non-T=1 shells.

### Encapsulin pore engineering yields polymorphic protein shells with high cargo capacity

To further characterize Mx_pmut shells in the absence or presence of SNAP-tag cargo, we carried out single particle cryo-EM analysis (Supplementary Fig. 3 and 4). We found that Mx_pmut without cargo formed T=1 shells while the cargo-loaded sample contained two distinct shell assemblies of T=3 (35 nm, 180 protomers) and, surprisingly, T=4 (39 nm, 240 protomers) icosahedral symmetry, determined to 3.14 and 3.46 Å resolution, respectively (Fig. 2a,b). This polymorphic behavior of cargo loaded Mx_pmut shells explains the broad shell size distribution observed via negative stain TEM analysis (Fig. 1f). The asymmetric unit of T=3 shells contained three unique protomers (A-C) while the T=4 shell contained four (A-D) (Fig. 2c). T=3 shells constituted ca. 75% of the sample while T=4 shells comprised the remaining ca. 25%. Both T=3 and T=4 shells contained strong density for the Mx-targeting peptide which allowed for confident atomic model building of the nine strongly interacting TP residues (KRLTVGSLR) and confirmed close to complete SNAP-tag cargo loading with T=3 shells containing ca. 180 and T=4 shells ca. 240 copies of SNAP-tag cargo (Fig. 2d). This is the first instance of T=3/T=4 structural polymorphism reported for encapsulins and overall, only the second report of a T=4 encapsulin shell^45^. As only T=1 shells were observed in the absence of cargo, cargo-shell co-assembly appears to be crucial for T=3/T=4 shell formation.

**Fig 2.**
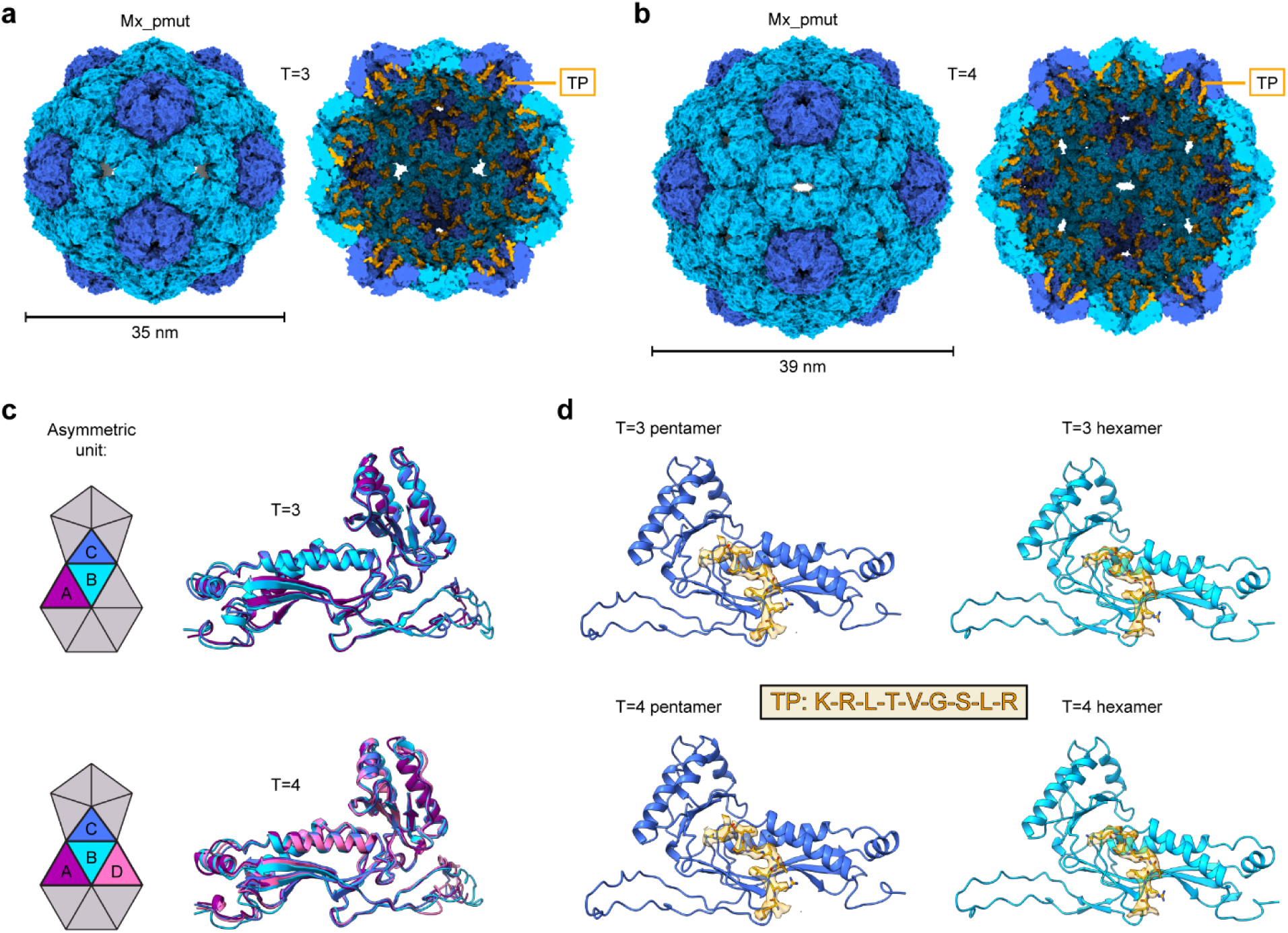
Structural characterization of polymorphic pore-engineered Mx_pmut encapsulins. **a**, Exterior (left) and cut-away (right) views of the T=3 Mx_pmut shell. Targeting peptides are shown in orange. **b**, Exterior (left) and cut-away (right) views of the T=4 Mx_pmut shell. Targeting peptides are shown in orange. **c**, Asymmetric unit (ASU) diagram for T=3 and T=4 Mx_pmut shells with unique protomers highlighted in different colors (left). Structural alignment of the unique protomers found in the T=3 and T=4 ASU. **d**, Targeting peptide (TP) densities and models (yellow) identified in the pentameric and hexameric facets of T=3 and T=4 Mx_pmut shells. The nine TP residues with strong density that could be confidently modeled are shown (KRLTVGSLR).

### Mx_pmut shells exhibit expanded rigid pores with favorable properties

Both T=3 and T=4 Mx_pmut shells contain pores of substantially increased size compared to Mx_wt. Pore-engineered T=3 shells exhibit five-fold pores of 12.6 Å (97% increase over Mx_wt) and three-fold pores of up to 21.6 Å (500% increase) as measured based on the atomic shell model (Fig. 3a,b). HOLE simulations resulted in even more pronounced percentage increases in pore diameters for Mx_pmut of 182% (five-fold pore: 3.4 Å vs 9.6 Å) and 1,413% (three-fold pore: 0.8 Å vs 12.1 Å). Similar results and pore size increases were obtained for pore-engineered T=4 shells (Fig. 3c,d). By removing the α6-α7 loop in Mx_pmut protomers, and thus the prior pore residues E_197_ and K_198_, a less charged pore exterior was realized with the pore interior and pore transit remaining slightly negatively charged (Fig. 3). In contrast to previously reported encapsulin pore mutants, all Mx_pmut pores in T=3 and T=4 shells exhibited well defined cryo-EM pore densities as supported by local resolution analysis (Supplementary Fig. 5). Flexible or partially disordered pores would likely negatively affect molecular flux across the protein shell. Thus, Mx_pmut shells may represent a promising system for challenging nanoreactor applications.

**Fig 3.**
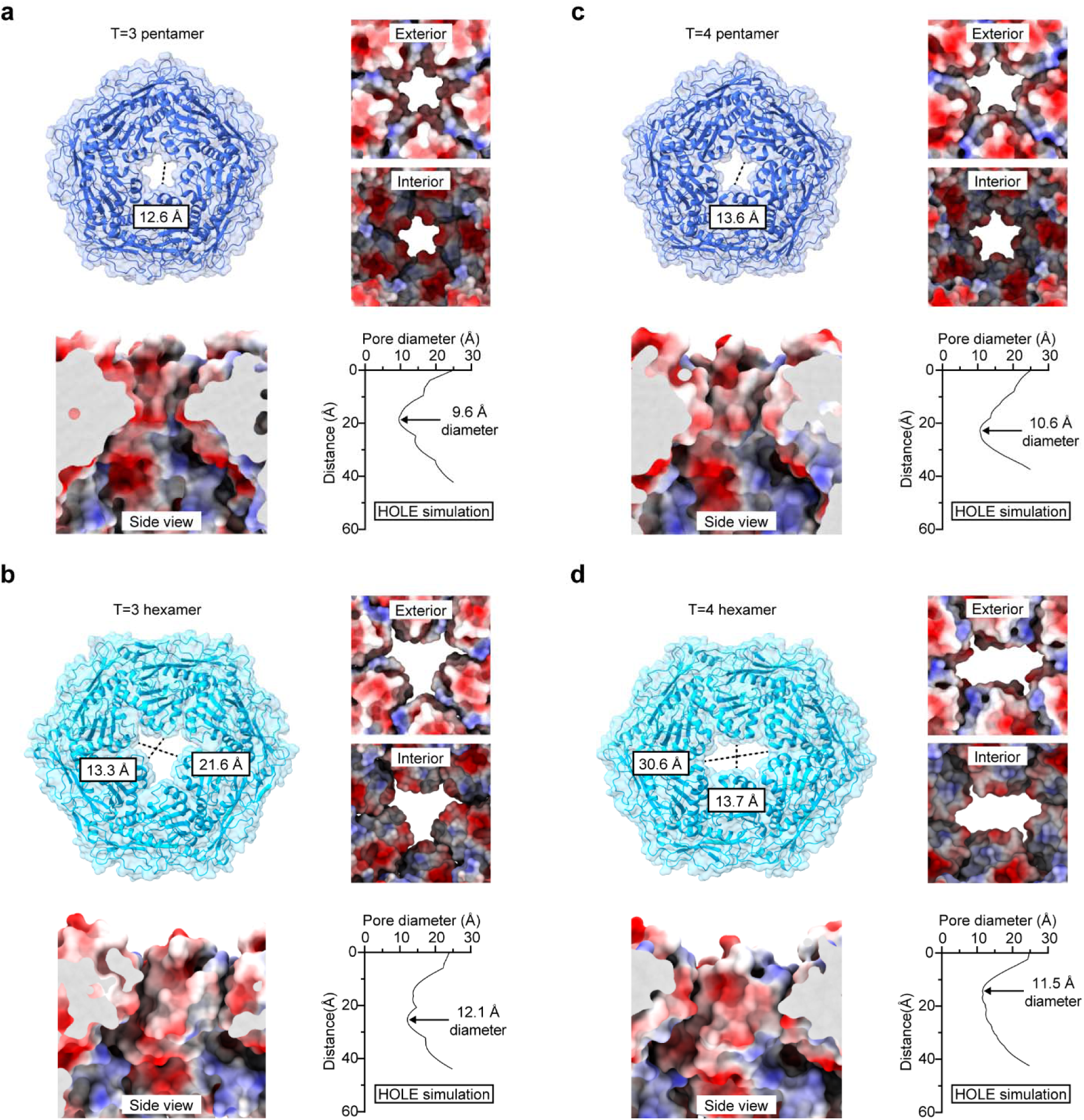
Pore analysis of engineered T=3 and T=4 Mx_pmut encapsulin shells. **a**, Analysis of the five-fold pore of the T=3 Mx_pmut shell. Top left: Exterior view down the five-fold axis of symmetry onto the five-fold pore of Mx_pmut. Pore diameter as measured based on the atomic model is shown. Top right: Close-up exterior and interior view of the five-fold pore in electrostatic surface representation. Bottom left: Side view perpendicular to the five-fold symmetry axis highlighting the electrostatic properties of the pore channel. Bottom right: Pore diameter profile as derived from HOLE simulations highlighting the narrowest point of the pore. **b**, Analysis of the three-fold pore of the T=3 Mx_pmut shell. Top left: Exterior view down the three-fold axis of symmetry onto the three-fold pore of Mx_pmut. Pore diameter as measured based on the atomic model is shown. Top right: Close-up exterior and interior view of the three-fold pore in electrostatic surface representation. Bottom left: Side view perpendicular to the three-fold symmetry axis highlighting the electrostatic properties of the pore channel. Bottom right: Pore diameter profile as derived from HOLE simulations highlighting the narrowest point of the pore. **c**, Analysis of the five-fold pore of the T=4 Mx_pmut shell. Top left: Exterior view down the five-fold axis of symmetry onto the five-fold pore of Mx_pmut. Pore diameter as measured based on the atomic model is shown. Top right: Close-up exterior and interior view of the five-fold pore in electrostatic surface representation. Bottom left: Side view perpendicular to the five-fold symmetry axis highlighting the electrostatic properties of the pore channel. Bottom right: Pore diameter profile as derived from HOLE simulations highlighting the narrowest point of the pore. **d**, Analysis of the three-fold pore of the T=4 Mx_pmut shell. Top left: Exterior view down the three-fold axis of symmetry onto the three-fold pore of Mx_pmut. Pore diameter as measured based on the atomic model is shown. Top right: Close-up exterior and interior view of the three-fold pore in electrostatic surface representation. Bottom left: Side view perpendicular to the three-fold symmetry axis highlighting the electrostatic properties of the pore channel. Bottom right: Pore diameter profile as derived from HOLE simulations highlighting the narrowest point of the pore.

### Increased pore size improves molecular flux across protein shells

We next set out to investigate how the increased pore size of Mx_pmut shells would affect the molecular flux of small molecules into the protein compartment interior. We devised an assay to monitor molecular flux in real time, utilizing a model system encapsulating the self-labeling protein SNAP-tag inside Mx_pmut shells. After addition of fluorescent SNAP-tag substrates, covalent modification of encapsulated SNAP-tag is used as a measure of molecular flux into the Mx_pmut interior. SNAP-tag labelling can be detected in real time by following the fluorescence polarization (FP) signal of the sample (Fig. 4a). Upon covalent binding of fluorescent SNAP-tag substrates to free and especially encapsulated SNAP-tag, an increase in the FP signal is expected as fluorophores are now stably bound to a protein (free SNAP-tag) or megadalton protein complex (encapsulated SNAP-tag), thus severely decreasing rotational motion compared to unbound fluorophores (Fig. 4a,b).

**Fig 4.**
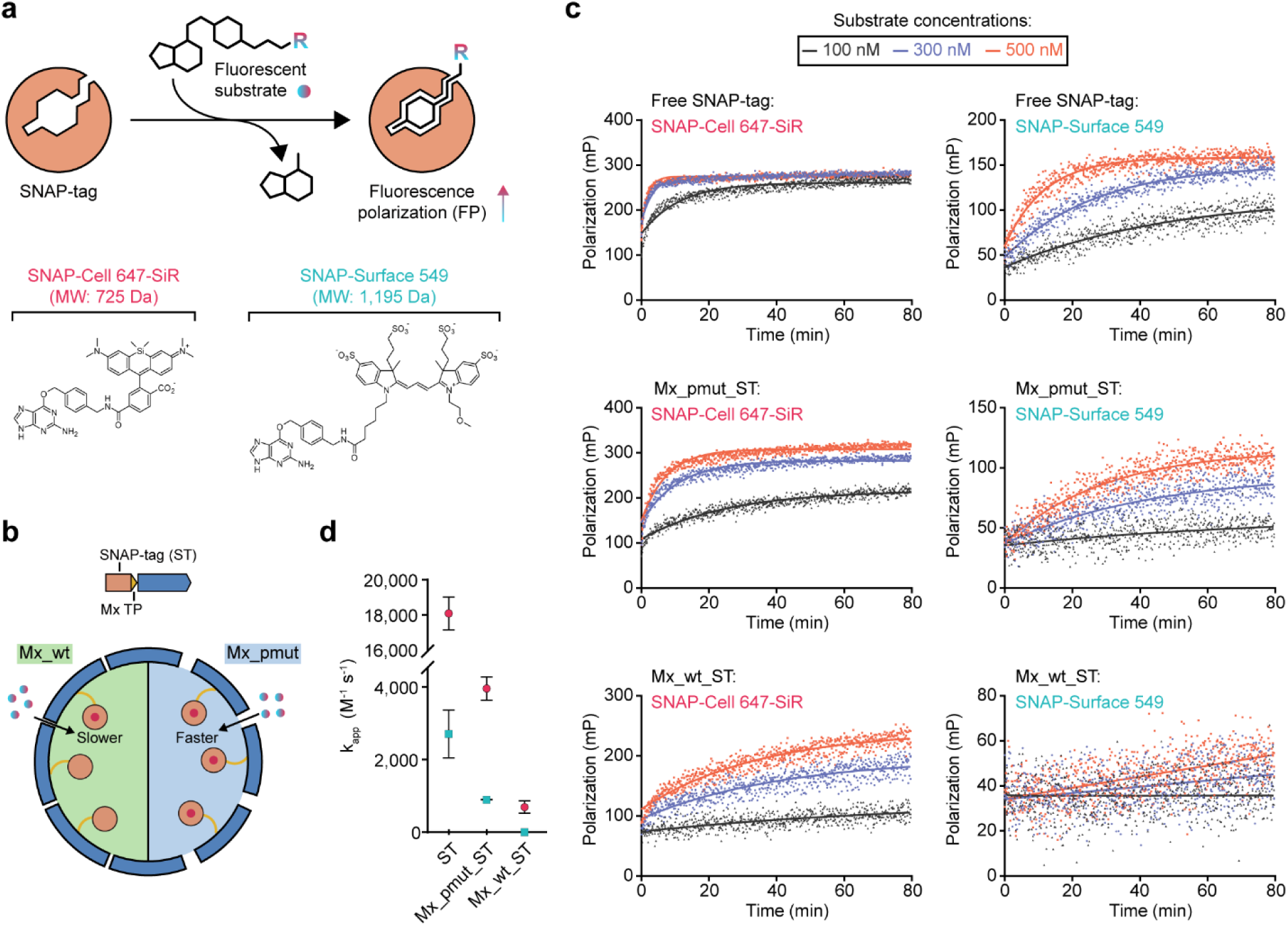
Molecular flux analysis and comparison of Mx_pmut and Mx_wt encapsulin shells. **a**, SNAP-tag-based fluorescence polarization assay scheme (top). The two SNAP-tag labeling substrates used (bottom). **b**, Schematic illustrating differential molecular flux of SNAP-tag substrates through the Mx_wt and Mx_pmut shells. **c**, Fluorescence polarization time courses at three different substrate concentrations for free SNAP-tag, SNAP-tag encapsulated in Mx_pmut shells (Mx_pmut_ST), and SNAP-tag encapsulated in Mx_wt shells (Mx_wt_ST) fit to a single-step pseudo-first-order kinetic model. Assays were carried out in triplicate. The substrate used is indicated in each plot. **d**, Apparent rate constants (k_app_) derived from the fluorescence polarization assays. Standard errors of three independent experiments are shown.

For our FP assays, we used two different fluorescent SNAP-tag substrates – SNAP-Cell 647-SiR (725 Da) and SNAP-Surface 549 (1,195 Da) – and monitored their labelling time courses with free SNAP-tag (ST), SNAP-tag encapsulated inside Mx_wt (Mx_wt_ST), and SNAP-tag encapsulated inside Mx_pmut (Mx_pmut_ST) (Fig. 4c). We selected the abovementioned substrates due to their different molecular weights and sizes (Fig. 4a), which represent the upper molecular weight and size ranges of common small molecule enzyme substrates and cofactors (ca. 500 to 1,000 Da). The primary goal of our FP assays is the comparison between the molecular flux behavior of Mx_wt and pore-engineered Mx_pmut. For both substrates, the labeling reaction occurred fastest for free SNAP-tag, followed by Mx_pmut_ST, and then Mx_wt_ST. This highlights that protein shells in general represent a diffusion barrier for the substrate and decrease labeling efficiency. When comparing the Mx_pmut_ST and Mx_wt_ST reactions, increasing pores size does have a marked effect on the observed molecular flux behavior. Specifically, for SNAP-Cell 647-SiR, the smaller substrate, increasing pore size allows faster molecular flux across the shell when compared to wild-type shells while for the larger SNAP-Surface 549 substrate, larger pores appear to expand the upper size threshold of substrates able to transit the shell with the larger substrate mostly excluded from the shell interior of Mx_wt. Apparent labeling rate constants (k_app_) derived for the different labeling reactions confirmed this trend. The pore-engineered Mx_pmut system showed substantially increased k_app_ values (3,950 M^-1^ s^-1^, 464% increase) when compared to Mx_wt (700 M^-1^ s-1) for the smaller SNAP-Cell 647-SiR substrate. No k_app_ could be calculated for Mx_wt in reactions using the larger SNAP-Surface 549 substrate, confirming that Mx_wt pores are too small to allow flux of this substrate. Mx_pmut on the other hand allowed the derivation of a k_app_ value (900 M^-1^ s^-1^), even for the large SNAP-Surface 549 substrate confirming that the effective size threshold for small molecule flux across the protein shell was increased. Interestingly, in labeling reactions using SNAP-Cell 647-SiR, the FP signal of saturated Mx_pmut_ST was higher than that of saturated free SNAP-tag. This is likely due to the large difference in size and molecular weight between the labeled protein components. Unlike free SNAP-tag, encapsulated SNAP-tag is stably anchored to a megadalton protein shell likely resulting in substantially decreased rotational motion when compared to the 19 kDa free SNAP-tag, thus resulting in an increased FP signal.

### Improved performance of pore-engineered enzyme nanoreactors

To investigate the effect of improved molecular flux on enzyme nanoreactor performance, we designed two prototype nanoreactor systems based on well characterized enzymes for both the pore-engineered Mx_pmut and native Mx_wt. Initially, the robust luciferase enzyme NanoLuc was selected which is able to carry out the cofactor-independent decarboxylation of the relatively large substrate fluorofurimazine (432 Da) with the concomitantly generated blue light (460 nm) serving as a convenient luminescence readout (Fig. 5a). After expression and purification, we were able to confirm that NanoLuc tagged with the Mx targeting peptide could be successfully encapsulated inside both Mx_pmut and Mx_wt with comparable cargo loading efficiencies of 70% (Fig. 5b and Supplementary Fig. 1). Again, a mixture of T=3 and T=4 shells could be observed (Supplementary Fig. 2). Subsequent NanoLuc nanoreactor activity assays showed that the turnover number for the Mx_pmut_NanoLuc nanoreactor was substantially improved (130% increase) when compared to the Mx_wt_NanoLuc design highlighting that increased pore size and improved molecular flux has a marked effect on the activity of encapsulated NanoLuc when using a large substrate (Fig. 5c). Next, we designed a set of nanoreactors encapsulating the well characterized promiscuous *Pyrococcus furiosus* alcohol dehydrogenase D (AdhD) which utilizes the universal and large redox co-substrate NADH (665 Da) to reduce diverse carbonyl groups to the corresponding alcohols. AdhD-loaded T=3 and T=4 Mx_pmut– and Mx_wt-based nanoreactors could be successfully produced and exhibited high and comparable cargo loading efficiencies of 98% (Fig. 5d and Supplementary Fig. 2). AdhD nanoreactor assays were carried out using dihydroxyacetone phosphate and NADH as substrates and showed that pore-engineered Mx_pmut nanoreactors displayed a turnover number increase of 89% when compared to Mx_wt-based nanoreactors.

**Fig 5.**
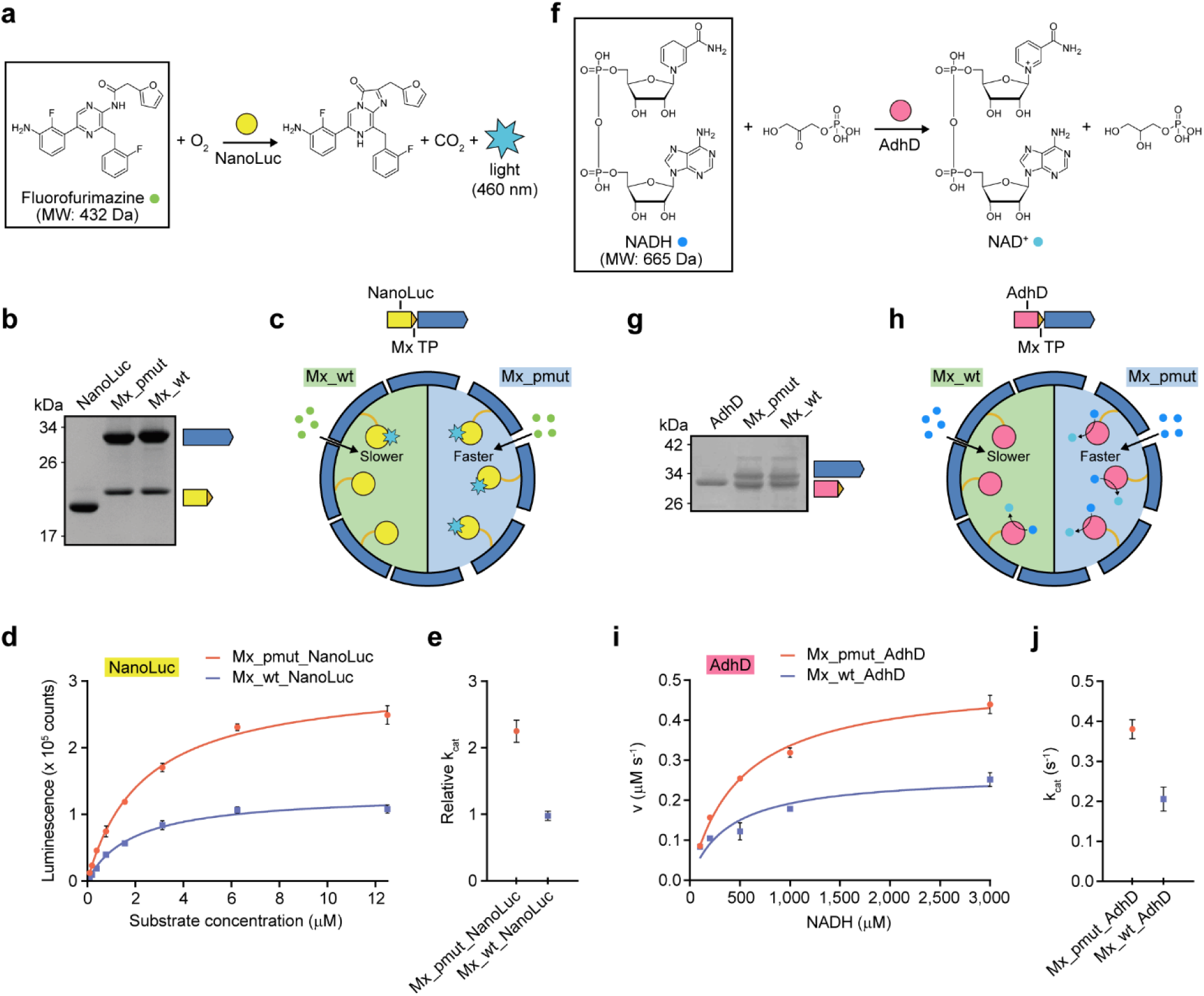
Characterization and comparison of Mx_pmut– and Mx_wt-based nanoreactors. **a**, Scheme of the NanoLuc-catalyzed reaction highlighting fluorofurimazine as the largest substrate involved in NanoLuc catalysis. **b**, SDS-PAGE analysis of free NanoLuc and Mx_pmut– and Mx_wt-encapsulated NanoLuc highlighting comparable cargo loading. **c**, Schematic illustrating differential molecular flux of the NanoLuc substrate fluorofurimazine through the Mx_wt and Mx_pmut shells. **d**, Substrate concentration-dependent activity of NanoLuc encapsulated inside Mx_pmut and Mx_wt shells. Error bars represent standard error of three independent experiments. **e**, Comparison of relative turnover numbers (k_cat_) for Mx_pmut– and Mx_wt-based NanoLuc nanoreactors. Error bars represent standard error of three independent experiments. **f**, Scheme of the AdhD-catalyzed reaction highlighting NADH as the largest substrate involved in AdhD catalysis. **g**, SDS-PAGE analysis of free AdhD and Mx_pmut– and Mx_wt-encapsulated AdhD highlighting comparable cargo loading. **h**, Schematic illustrating differential molecular flux of NADH through the Mx_wt and Mx_pmut shells. **i**, Saturation kinetics analysis of Mx_pmut– and Mx_wt-based AdhD nanoreactors. Error bars represent standard error of three independent experiments. **j**, Comparison of turnover numbers (k_cat_) for Mx_pmut– and Mx_wt-based AdhD nanoreactors. Error bars represent standard error of three independent experiments.

One of the primary benefits of enzyme encapsulation is the improved stability of sequestered enzymes and their increased protection from proteolytic degradation and other harsh conditions, thus improving overall enzyme activity and longevity in challenging environments. To investigate if pore engineering has any negative impact on shell integrity and stability, we carried out dynamic light scattering analyses coupled with thermal ramps on both Mx_pmut– and Mx_wt-based enzyme nanoreactors. No significant differences in terms of polydispersity or stability between Mx_pmut and Mx_wt shells encapsulating the same cargo could be detected (Supplementary Fig. 6). This suggests that pore engineering has little effect on shell integrity as long as the primary interaction interfaces between protomers are maintained. Next, the protective function of the Mx_pmut and Mx_wt shells was assayed via incubations with the protease trypsin and freshly prepared bacterial lysate. The activity of encapsulated NanoLuc – in both Mx_pmut and Mx_wt – was found to be substantially less affected than that of free NanoLuc after 6 h of tryptic digest or overnight lysate exposure while little difference in AdhD activity after similar trypsin incubation could be observed between the encapsulated or free samples (Supplementary Fig. 7). These results suggest that encapsulation can increase the protease resistance of proteins susceptible to proteolytic digest and that this protective capacity is not negatively affected by pore engineering.

Our results highlight the potential of pore-engineering as a general strategy to address nanoreactor performance issues caused by suboptimal shell porosity without compromising protein shell assembly or stability. In addition, the encapsulin-based Mx_pmut shell with its modular and efficient cargo loading modality and optimized shell porosity has the potential to be used as a versatile nanoreactor platform in combination with a broad range of biomedically or biotechnologically relevant enzymes, independent of small molecule substrate or cofactor size.

### Co-encapsulation of multiple enzymes inside Mx_pmut enables *in situ* cofactor recycling

One powerful application of enzyme nanoreactors is the co-localization of multiple distinct enzymes to build artificial metabolons^4^. One aspect of naturally occurring metabolons is an intrinsic cofactor recycling system, as for example realized in bacterial microcompartment-based ethanolamine or 1,2-propanediol utilization metabolons where the redox cofactor NAD^+^ is recycled inside the protein shell by a dedicated enzyme component^8^. Realizing such a system would substantially expand the potential use cases of engineered enzyme nanoreactors. To explore if such a cofactor recycling system could be implemented in Mx_pmut– and Mx_wt-based nanoreactors, specifically for the ubiquitous redox cofactor NADH, we designed nanoreactors for the co-encapsulation of NADH-dependent AdhD and NADH-generating *Stutzerimonas stutzeri* phosphite dehydrogenase (PTDH). PTDH is able to oxidize inorganic phosphite to inorganic phosphate whilst reducing NAD^+^ to NADH. In a two-enzyme AdhD-PTDH nanoreactor, the initially added small amount of NADH would first be consumed by AdhD and converted to NAD^+^ which in turn would then be utilized by PTDH and regenerated to NADH, thus closing the recycling loop (Fig. 6a). We were able to successfully produce and purify AdhD– and PTDH-loaded Mx_pmut and Mx_wt nanoreactors which exhibited identical loading ratios of 85% AdhD and 15% PTDH (Fig. 6b). We now separately tested PTDH and AdhD activity in the context of the Mx_pmut and Mx_wt two-enzyme nanoreactors. PTDH was active in both co-encapsulation systems and showed higher turnover (84% increase) in the Mx_pmut-based nanoreactor (Supplementary Fig. 8) while AdhD activity was not negatively affected by co-encapsulation when compared to AdhD-only nanoreactors (Supplementary Fig. 9). We subsequently carried out cofactor recycling assays by first adding excess amounts of AdhD substrate (dihydroxyacetone phosphate, 10 mM) and small amounts of NADH (130 μM) to nanoreactor samples. After NADH was fully oxidized to NAD^+^, an excess of PTDH substrate (inorganic phosphite, 10 mM) was added and NADH regeneration was followed by monitoring the fluorescence increase at 465 nm. We observed NADH regeneration in both Mx_pmut and Mx_wt nanoreactors until equilibrium between NADH production and consumption was reached. Notably, NADH regeneration was faster and the equilibrium concentration of NADH was higher in Mx_pmut when compared to Mx_wt, likely caused by improved molecular flux and the improved PTDH activity previously observed for Mx_pmut-based nanoreactors. Our results highlight the usefulness of encapsulin-based nanoreactors for the construction of more complex multi-enzyme nanoreactor systems and the importance of pore size on overall nanoreactor performance.

**Fig 6.**
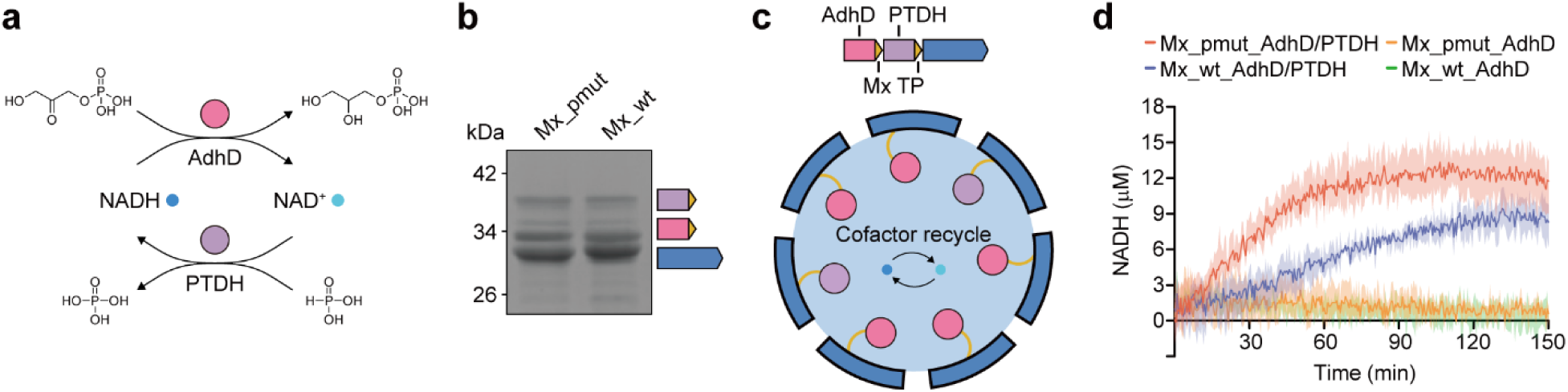
Characterization of multi-enzyme Mx_pmut nanoreactors for *in situ* cofactor recycling. **a**, Reaction scheme for NADH recycling using AdhD and PTDH. **b**, SDS-PAGE gel of AdhD and PTDH co-encapsulation nanoreactors highlighting comparable relative cargo loading of Mx_pmut and Mx_wt. **c**, Schematic illustrating Mx_pmut-based AdhD– and PTDH-loaded nanoreactors for NADH recycling. **d**, Time course of NADH regeneration after the addition of phosphite to NADH-depleted samples.

## Discussion

In this study, we design and characterize a pore-engineered encapsulin shell (Mx_pmut) and demonstrate its improved small molecule molecular flux behavior. We show that Mx_pmut-based enzyme nanoreactors exhibit improved catalytic performance while maintaining the ability to stabilize and protect encapsulated enzymes.

Previous attempts at encapsulin pore engineering have been restricted to small T=1 encapsulin shells with limited cargo loading capacity^38–40^. As larger T=3 and T=4 shells contain two distinct types of pentameric and hexameric pores, both formed by a single type of shell protomer, any protomer mutations have to maintain both of these interactions, otherwise, no T=3 or T=4 shell can be assembled. Beyond their small size, previously reported pore engineered T=1 encapsulin systems often suffered from reduced solubility and flexible, often poorly defined, pores, impeding small molecule molecular flux across the shell^40^.

In response to these challenges, we have established the first non-T=1 pore-engineered encapsulin system – Mx_pmut – based on the model T=3 encapsulin from *M. xanthus* (Mx_wt). Mx_pmut forms T=1 shells in the absence of cargo while cargo protein co-expression results in highly cargo-loaded polymorphic T=3 and T=4 shells. This highlights that the structural integrity of hexameric facets can be easily compromised by pore mutations and supports our hypothesis that cargo co-expression can stabilize hexameric facets in pore-engineered systems. At the same time, the requirement of cargo-shell co-assembly for T=3/T=4 shell formation guarantees high cargo loading with substantial loading percentages (70 to 98%) observed across various cargo types in this study. Further, our structural analysis of Mx_pmut highlighted well-defined substantially increased pores compared to Mx_wt, closely resembling our original design.

Using SNAP-tag-based fluorescence polarization assays as well as a number of single– and two-enzyme proof-of-concept nanoreactor designs relying on large substrates, we show that the expanded pore size of Mx_pmut markedly increases molecular flux and extends the upper size limit for small molecules able to efficiently cross the shell. This will allow Mx_pmut to be used as a nanoreactor platform in combination with a wide array of enzymes, particularly those utilizing large substrates or cofactors, which was previously hard to realize in encapsulin-based systems due to small native pores. The consistently observed improved performance of our Mx_pmut-based test enzyme nanoreactors underscores the significance of pore size in determining overall nanoreactor performance.

One of the potential concerns with increasing pore size is the loss of structural integrity and protective capacity of the shell. It has been shown that large pores (>3 nm) can allow small proteins, like RNase A (14 kDa), to access the luminal space within protein cages, thereby negating the protective effect of cargo encapsulation^55^. As many small proteases have been identified with molecular weights between 5 and 20 kDa^56,57^, large pores could lead to substantial challenges with respect to proteinaceous cargo stability in certain nanoreactor applications. The engineered pores in Mx_pmut are generally below the threshold of 3 nm, making it likely that cargo will still be protected from nearly all hydrolytic enzymes.

While this study establishes Mx_pmut as a general nanoreactor platform technology, certain aspects of designing nanoreactors go beyond an optimized shell – achieved in this study – and an efficient cargo loading mechanism – inherently found in encapsulin systems. Generally, the encapsulation of enzymes within protein nanocages creates a crowded and confined environment that can influence the catalytic activity of encapsulated enzymes, independent of substrate-or cofactor-related issues^58^. Enzymes reliant on multimerization for activity may benefit from this self-crowding effect, whereas enzymes requiring substantial conformational changes for activity could be negatively affected by encapsulation. In the context of our Mx_pmut_AdhD/PTDH two-enzyme nanoreactor, the activity of PTDH significantly decreased upon encapsulation, possibly due to unfavorable interactions within the shell inhibiting the conformational changes required by PTDH to be fully active^59^. Selecting enzymes that do not suffer from inherent activity loss upon encapsulation will be crucial to fully exploit the benefit of Mx_pmut-based nanoreactors.

In sum, through pore engineering, we have built a modular encapsulin-based enzyme nanoreactor platform – Mx_pmut. We demonstrate that rational pore engineering can be a successful strategy for enhancing the overall performance of protein-based enzyme nanoreactors. Our work lays the groundwork for future application-focused studies aimed at utilizing advanced enzyme nanoreactors in biomedical and biotechnological contexts.

## Methods

### Molecular biology and cloning

All constructs used in this study, except Mx_pmut, were initially ordered from Integrated DNA Technologies (IDT) as *E. coli* codon-optimized gBlocks and their expression optimized, if necessary, by introducing mutations at the 5’ end of the respective genes, yielding the final expression constructs (Supplementary Table 1). The Mx_pmut construct was obtained through overhang PCR using the Mx_wt gene as a template (Supplementary Table 2). All genes were cloned into the pETDuet-1 vector. Genes encoding cargo proteins were inserted into multiple cloning site 1 (MCS1) and encapsulin genes were inserted into MCS2. The respective vector backbones for different constructs were amplified via PCR and assembled with inserts carrying appropriate overhangs via Gibson assembly. In case of the AdhD and PTDH co-encapsulation system, the AdhD and PTDH genes were inserted into MCS1 as a two-gene operon, separated by a short intergenic sequence containing the same ribosome binding site (RBS) as found in MCS1 and MCS2. *E. coli* BL21 (DE3) cells were transformed with the assembled plasmids via electroporation and were confirmed through Sanger sequencing (Eurofins Scientific).

### Protein Expression and purification

All expression experiments were carried out using lysogeny broth (LB) medium supplemented with 100 mg/mL ampicillin. 500 mL of fresh LB medium were inoculated 1:100 using a 5 mL overnight culture, grown at 37°C to an OD600 of 0.5-0.6, and then induced with 0.1 mM isopropyl β-D-1-thiogalactopyranoside (IPTG). After induction, cultures were grown at 30°C overnight for ca. 18 h and harvested via centrifugation (5,000 g, 12 min, 4°C). The resulting cell pellets were frozen and stored at –80°C until further use.

Frozen cell pellets were resuspended in 5 mL/g (wet cell mass) of Tris buffer (20 mM Tris, 150 mM NaCl, pH 7.5). Lysis components [lysozyme (0.5 mg/mL), Benzonase® nuclease (25 units/mL), MgCl_2_ (1.5 mM), SIGMAFAST EDTA-free protease inhibitor cocktail (one tablet per 100 mL)] were added and cells were incubated on ice for 20 min. Samples were then sonicated at 65% amplitude and a pulse time of 10 s on and 20 s off for 5 min total (Model 120 Sonic Dismembrator, Fisher Scientific). After sonication, samples were clarified by centrifugation (10,000 g, 15 min, 4°C). To the supernatant, NaCl and PEG-8000 were added to a final concentration of 0.5 M and 10%, respectively, and incubated on ice for 40 min, followed by centrifugation (8,000 g, 10 min, 4°C). The supernatant was removed, and the pellet was resuspended in 4 mL Tris buffer (pH 7.5) and filtered using a 0.2 μm syringe filter. The filtered sample was subjected to size exclusion chromatography (SEC) using a Sephacryl S-500 16/60 column and Tris buffer (pH 7.5) at a flow rate of 1 mL/min. Fractions were evaluated using SDS-PAGE and encapsulin-containing fractions were combined, concentrated, and dialyzed using Amicon filter units (100 kDa MWCO) and Tris buffer without NaCl (20 mM Tris, pH 7.5). The low salt sample was then loaded on a HiPrep DEAE FF 16/10 Ion Exchange column at a flow rate of 3 mL/min to remove nucleic acid contamination. Encapsulin-containing fractions were concentrated, centrifuged (10,000 g, 10 min, 4°C), and then subjected to another round of SEC using a Superose 6 10/300 GL column and Tris buffer (pH 7.5) at a flow rate of 0.5 mL/min. Purified proteins were stored in Tris buffer (pH 7.5) at 4°C until further use.

### SDS polyacrylamide gel electrophoresis

SDS polyacrylamide gel electrophoresis (SDS-PAGE) was conducted in an Invitrogen XCell SureLock system using Novex 14% Tris-Glycine Mini Protein Gels with SDS running buffer. 4-5 μg of each sample were mixed with 4X SDS sample buffer, heated for 4 min at 98°C, briefly centrifuged, and subsequently loaded onto the gel. SDS-PAGE analysis was conducted at a constant voltage of 225 V for 42 min at room temperature. The Spectra Multicolor Broad Range Protein Ladder was used as a molecular weight marker.

### Negative stain transmission electron microscopy (TEM)

Encapsulin samples for negative-stain TEM were diluted to 0.15 mg/mL in Tris buffer (pH 7.5). Gold grids (200-mesh coated with Formvar-carbon film, EMS #FCF200-Au-EC) were made hydrophilic by glow discharge at 5 mA for 60 s (easiGlow, PELCO). 4 μL of sample was added to the grid and incubated for 1 min, wicked with filter paper, and washed with 0.75% uranyl formate before staining with 0.75% uranyl formate for 1 min. Stain was removed using filter paper and the grid was dried for at least 20 min before imaging. TEM micrographs were captured using a Morgagni transmission electron microscope at 100 kV at the University of Michigan Life Sciences Institute.

### Pore dimension measurements

The HOLE package (2.2.005) was used for simulations to determine the radius of five– and three-fold encapsulin pores following standard protocols^60^. The default file was used to determine atomic van der Waals radii. Pore measurements were conducted with a cutoff radius of 20 Å.

### Dynamic light scattering (DLS) analysis

All sizing and polydispersity measurements were carried out on an Uncle instrument (Unchained Labs) at 15°C in triplicate. All encapsulin samples were adjusted to 0.4 mg/mL of monomer using Tris buffer (pH 7.5), centrifuged (10,000 g, 10 min, 4°C), and then immediately analyzed via DLS.

### Cryo-electron microscopy (cryo-EM)

*Sample preparation:* Purified samples of Mx_pmut T=1 and Mx_pmut T=3/T=4 were concentrated to 3 mg/mL and 4.1 mg/mL, respectively, in 150 mM NaCl, 25 mM Tris pH 8.0. 3.5 µL of protein samples were applied to freshly glow discharged grids (Quantifoil R1.2/1.3 Cu 200 mesh) and prepared by plunge freezing in liquid ethane using an FEI Vitrobot Mark IV (100% humidity, 22°C, blot force 20, blot time 4 s, wait time 0 s). The grids were immediately clipped and stored in liquid nitrogen until data collection.

*Data collection:* Cryo-EM movies for Mx_pmut T=1 and Mx_pmut T=3/T=4 were collected using a ThermoFisher Scientific Talos Arctica cryo-electron microscope operating at 200 kV equipped with a Gatan K2 direct electron detector. 711 movies for Mx_pmut T=1 and 706 movies for Mx_pmut T=3/T=4 were collected from a single grid for each sample using the Leginon^61^ software package at a magnification of 45,000x, pixel size of 0.91 Å, defocus range of –1.0 µm to –1.8 µm, exposure times of 4 or 6 s, frame time of 200 ms, and a total dose of 39.18 e^-^/Å^2^ for Mx_pmut T=1 and 39.56 e^-^/Å^2^ for Mx_pmut T=3/T=4.

*Data processing:* CryoSPARC 3.10.0^62^ was used to process the Mx_pmut T=1 dataset. 711 movies were imported, motion corrected by patch motion correction, and the CTF fit was estimated using patch CTF estimation. Exposures with CTF fit resolutions worse than 6 Å were discarded from the dataset, resulting in 692 remaining movies. 186 particles were selected manually and used to create templates for particle picking. Template picker was then used to select particles and 66,813 particles were extracted from the dataset with a box size of 360 pixels. The particles were then sorted by two rounds of 2D classification, resulting in 62,306 remaining particles. An initial volume was created by ab-initio reconstruction using two classes and I symmetry, resulting in a majority class containing 61,808 particles. These particles were then used for homogeneous refinement against the ab-initio map with I symmetry imposed, per-particle defocus optimization, per-group CTF parameterization, and Ewald sphere correction enabled, resulting in a 2.86 Å map.

CryoSPARC 4.10.0^62^ was used to process the Mx_pmut T=3/T=4 datasets. 706 movies were imported, motion corrected using patch motion correction, and the CTF fit was estimated using patch CTF estimation. Movies with CTF fit resolutions worse than 6 Å were discarded from the dataset, resulting in 692 remaining movies. Manual picker was used to select 548 particles of Mx_pmut T=3 and Mx_pmut T=4, which were used to generate templates for particle picking. 37,600 particles containing both T=3 and T=4 particles were identified using template picker, extracted with an initial box size of 560 pixels, which were then downsampled to a box size of 256 pixels for downstream data processing. Two rounds of 2D classification were used to sort Mx_pmut T=3/T=4 particles, resulting in 12,992 particles. An initial volume containing 12,967 particles was created using ab-initio reconstruction with two classes and I symmetry. These particles were then used for homogeneous refinement against the ab-initio map using I symmetry, per-particle defocus optimization, per-group CTF parametrization, and Ewald sphere correction enabled, resulting in a 3.14 Å map of Mx_pmut T=3. Two rounds of 2D classification were then used to select 4,394 particles of Mx_pmut T4. An initial volume containing 4,386 particles was created using ab-initio reconstruction with two classes and I symmetry. These particles were then used for homogeneous refinement against the Mx_pmut T4 ab-initio map using I symmetry, per-particle defocus optimization, per-group CTF parametrization, and Ewald sphere correction enabled, resulting in a final 3.46 Å resolution map of Mx_pmut T4.

*Model building:* For building the Mx_pmut T=1 shell model, a T=1 starting model of the *Myxococcus xanthus* Enc A protomer (PDB: 7S21)^46^ was placed manually into the map using ChimeraX v.1.2.5^63^, followed by improving the map fit using the fit-in-map command. The model was then manually modified to match the Mx_pmut T=1 amino acid sequence, and manually refined against the density map using Coot v0.9.8.1^64^. Phenix v 1.20.1-4487-000 was then used to further refine the model by real-space refinement with three macrocycles, minimization_global enabled, local_grid_search enabled, and adp refinement enabled. NCS operators were then identified from the map using map_symmetry and applied to the model using apply_ncs to generate the icosahedral shell. The NCS-expanded shell was then refined again using real-space refinement with three macrocycles, minimization_global enabled, local_grid_search enabled, adp refinement enabled, and NCS constraints enabled. The BIOMT operators were identified using the find_ncs command, and manually placed into the header of the .pdb file containing a single asymmetric unit (ASU) of the NCS-refined model (Supplementary Table 3). The same protocol was used to build the Mx_pmut T=3 and Mx_pmut T=4 shell models, except a T=3 icosahedral assembly of *Myxococcus xanthus* Enc A was used as the starting model for both shells (PDB: 7S4Q)^46^.

### Fluorescence polarization (FP) assays

Binding kinetics with SNAP-tag using the SNAP-tag substrates SNAP-Cell 647-SiR and SNAP-Surface 549 were carried out by recording FP over time using microplate readers (SNAP-Cell 647-SiR: Perkin Elmer Envision (Ex: 620 nm / Em: 688 nm), SNAP-Surface 549: BMG PHERAstar (Ex: 540 nm / Em: 590 nm)). Protein samples were prepared by diluting free SNAP-tag (ST), Mx_pmut_ST, and Mx_wt_ST with DTT (final concentration: 1mM)-containing phosphate-buffered saline (PBS) to a final concentration of either 100 nM, 300 nM, or 500 nM with respect to ST. The concentrations of ST in Mx_pmut_ST and Mx_wt_ST samples were determined through gel densitometry using SDS-PAGE gels, calibrated with a series of known concentrations of free ST run on the same gels. FP assays were initiated by adding either SNAP-Cell 647-SiR or SNAP-Surface 549 to the protein samples prepared in flat black bottom non-binding surface 96-well plates (Corning 3991). FP measurements were conducted in 20 second intervals for 80 min at room temperature with a final reaction volume of 100 μL. All assays were conducted in triplicate. For the negative control, final SNAP-tag substrate concentrations of 100 nM, 300 nM, and 500 nM were used with Mx_wt. Non-linear regression curve analysis with one phase association fits were conducted to derive apparent rate constants (k_app_) utilizing GraphPad Prism 9.

### NanoLuc enzymatic assays

NanoLuc activity was quantified by measuring luminescence immediately following the addition of its substrate, fluorofurimazine, using a Synergy H1 plate reader. Protein samples for the assays were prepared by diluting free NanoLuc, Mx_pmut_NanoLuc, and Mx_wt_NanoLuc with Tris buffer (pH 7.5) to a final concentration of 50 pM with respect to NanoLuc. NanoLuc activity assays were initiated by adding serially diluted fluorofurimazine to the protein samples prepared in flat white bottom non-binding surface 96-well plates (Corning 3990). The final concentrations of fluorofurimazine ranged from 0.0977 μM to 12.5 μM, doubling in concentration at each dilution step. Luminescence readings were obtained in triplicate at room temperature, with a final reaction volume of 100 μL. As a background control, the same reaction mixtures without protein were used and subtracted from the luminescence signal of each sample. Non-linear regression curve analysis using the Michaelis–Menten model was conducted on substrate course initial rates using GraphPad Prism 9.

### AdhD enzymatic assays

AdhD activity was quantified by tracking the oxidation of NADH at 340 nm using a Synergy H1 plate reader. Tris buffer (pH 7.5)-based reaction buffers were prepared by adding final concentrations of 10 mM dihydroxyacetone phosphate (DHAP), 1 mM polymyxin, and varying amounts of NADH (final concentrations: 100 μM, 200 μM, 500 μM, 1000 μM, and 3000 μM). Polymyxin was added to the reaction buffer in order to mitigate background NADH oxidation observed in Mx_pmut_AdhD and Mx_wt_AdhD samples resulting from small amounts of contaminating type II NADH-quinone oxidoreductase (NDH-2). The AdhD activity assays were initiated by adding protein samples of free AdhD, Mx_pmut_AdhD, and Mx_wt_AdhD to the 37°C pre-incubated reaction buffer prepared in black with flat clear bottom 96-well plates (Corning 3631). The final concentrations of the protein samples were set to 1.32 μM with respect to AdhD. The absorbance at 340 nm (A340) was measured in triplicate for total 80 min at 37°C with a final reaction volume of 100 μL. As a background control, the same reaction mixtures without protein were used and subtracted from the A340 of each sample. A standard curve for NADH-A340 was recorded to convert A340 signal to the corresponding concentration of NADH. Initial reaction rates of AdhD for varying concentrations of NADH were determined using the slope of the linear segment of the initial phase of the time course plots. Subsequently, non-linear regression curve analysis using the Michaelis–Menten model was conducted on substrate concentration-initial rate plots using GraphPad Prism 9.

### PTDH enzymatic assays

PTDH activity was quantified by tracking the reduction of NAD^+^ at 340 nm using a Synergy H1 plate reader. Tris buffer (pH 7.5)-based reaction buffers contained final concentrations of 10 mM phosphite, 1 mM polymyxin, and varying amount of NAD^+^ (final concentrations: 50 μM, 100 μM, 300 μM, 500 μM, and 1000 μM). The PTDH activity assays were initiated by adding protein samples of free PTDH, Mx_pmut_AdhD/PTDH, and Mx_wt_AdhD/PTDH to the 37°C pre-incubated reaction buffer prepared in black with flat clear bottom 96-well plates (Corning 3631). The final concentration of the protein samples were set to 1.41 μM with respect to PTDH. The A340 was measured in triplicate for a total of 150 min at 37°C with a final reaction volume of 100 μL. As background controls, the same reaction mixtures without protein were used and subtracted from the A340 signal of each sample. Initial reaction rates of PTDH for varying concentrations of NAD^+^ were determined using the slop of the linear segment of the initial phase of the time course. Subsequently, non-linear regression curve analysis using the Michaelis–Menten model was conducted using GraphPad Prism 9.

### AdhD/PTDH nanoreactor-based NADH regeneration

NADH regeneration assays using the AdhD/PTDH co-encapsulation system were quantified by tracking the characteristic fluorescence of NADH at 465 nm (E465) when excited at 340 nm using a Synergy H1 plate reader. NADH regeneration assays were carried out using Mx_pmut_AdhD/PTDH and Mx_wt_AdhD/PTDH along with Mx_pmut_AdhD, and Mx_wt_AdhD as negative controls. As the loading ratio of AdhD and PTDH in Mx_pmut_AdhD/PTDH and Mx_wt_AdhD/PTDH were identical, the final concentrations of all 4 nanoreactors were normalized to 1.5 μM with respect to AdhD. The reaction buffer used for initial NADH depletion consisted of 10 mM dihydroxyacetone phosphate, 1 mM polymyxin, and 130 μM NADH in Tris buffer (pH 7.5). The initial NADH depletion was started by adding protein sample to the 37°C pre-incubated reaction buffer prepared in triplicate with a final reaction volume of 80 μL. Right after adding the protein sample, 20 μL of reaction mixture were transferred into black flat bottom 384-well plates in duplicate and the E465 signal was measured over time at 37°C in order to tract NADH depletion. The remaining 40 μL of reaction mixture was incubated at 37°C until NADH was depleted. NADH depletion in each sample was accomplished by allowing the reaction to proceed until the E465 intensity reached a plateau, aligning with that of the control blank, which consisted of the corresponding reaction mixture without any NADH. Following the depletion of NADH in each sample, 15 μL of the reaction mixture was transferred into two separate 384-well plate wells of. Subsequently, 5 μL of TBS buffer (pH 7.5) was added to one of the aliquots as a background control, while an equal volume of a solution containing phosphite to achieve a final concentration of 10 mM was added to the other aliquot to initiate NADH regeneration. Upon phosphite/TBS buffer addition, E465 was measured for 150 min at 37°C. The amount of regenerated NADH over time was determined by subtracting the E465 values obtained from the background control from those of the samples. These corrected E465 values were then converted to NADH concentrations using an NADH standard curve.

### Enzyme stability assays

Trypsin incubation stability assays were carried out for NanoLuc– and AdhD-based nanoreactor systems. For NanoLuc-based system, purified free NanoLuc, Mx_pmut_NanoLuc, and Mx_wt_NanoLuc were diluted with Tris buffer (pH 7.5) to adjust NanoLuc concentration across samples to 3.4 μM. Subsequently, trypsin (Promega, USA) was added to each sample to achieve a NanoLuc to trypsin ratio of 20:1 (w/w) and was incubated at 37°C. Samples were takenafter 0, 2, 4, and 6 h post addition. At each designated time point, aliquots of the reaction mixture were frozen for later SDS-PAGE analysis. The remaining portions of the samples were used for the NanoLuc activity assay as described previously, employing a final concentration of 12.5 μM of the substrate fluorofurimazine, to assess NanoLuc activity after trypsin exposure. The same protocol was applied to the AdhD-based nanoreactor system where the final concentration of AdhD across samples was adjusted to 18.6 μM for subsequent trypsin incubations. Samples were taken after 0 and 6 h and AdhD activity was assayed as described previously, employing a final concentration of 3 mM of NADH substrate.

Bacterial cell lysate-based stability assay was conducted for NanoLuc-based nanoreactors. Cell lysate was prepared from *E. coli* Nissle 1917 as it possesses intact protease systems, in contrast to *E. coli* BL21 (DE3) which lacks most important cellular proteases. 500 mL of fresh LB medium was inoculated 1:100 using a 5 mL overnight *E. coli* Nissel 1917 culture and grown at 37°C overnight for ca. 18 h. Cells were harvested via centrifugation (5,000 g, 12 min, 4°C) and the resulting cell pellet was used to prepare cleared cell lysate as described previously. NanoLuc concentrations in free NanoLuc, Mx_pmut_NanoLuc, and Mx_wt_NanoLuc samples were normalized to 5 μM using Tris buffer (pH 7.5). Thereafter, 3 μL of each normalized sample was added to 100 μL of cleared cell lysate in triplicate and incubated at 37°C overnight for ca. 16 h. NanoLuc activity assays were then conducted as described previously, with a final concentration of 12.5 μM of the substrate fluorofurimazine.

## Data availability

Cryo-EM maps and structural models have been deposited in the Electron Microscopy Data Bank (EMDB) and the Protein Data Bank (PDB) and are publicly available. Mx_pmut T=1: EMDB-44383, PDB ID: 9B9I. Mx_pmut T=3: EMDB-44388, PDB ID: 9B9Q. Mx_pmut T=4: EMDB=44427, PDB ID: 9BC8.

## Acknowledgements

We acknowledge funding from the NIH (R35GM133325). Research reported in this publication was supported by the University of Michigan Cryo-EM Facility (U-M Cryo-EM). U-M Cryo-EM is grateful for support from the U-M Life Sciences Institute and the U-M Biosciences Initiative. Molecular graphics and analyses performed with UCSF ChimeraX, developed by the Resource for Biocomputing, Visualization, and Informatics at the University of California, San Francisco, with support from the National Institutes of Health R01GM129325 and the Office of Cyber Infrastructure and Computational Biology, National Institute of Allergy and Infectious Diseases.

## Author contributions

S.K. and T.W.G. designed the project. S.K. and M.P.A conducted the laboratory experiments, with S.K conducting all experiments except for cryo-EM data collection, data analysis, and model building. M.P.A. collected and analyzed the cryo-EM data. M.P.A. and T.W.G. processed the cryo-EM data. M.P.A. built the structural models. S.K and T.W.G. wrote the manuscript. T.W.G. oversaw the project in its entirety.

## References

1. Edwardson, T.G.W. et al. Protein Cages: From Fundamentals to Advanced Applications. Chem. Rev. 122, 9145–9197 (2022).

2. Cheah, L.C. et al. Artificial Self-assembling Nanocompartment for Organizing Metabolic Pathways in Yeast. ACS Synth. Biol. 10, 3251–3263 (2021).

3. Wang, M., Abad, D., Kickhoefer, V.A., Rome, L.H. & Mahendra, S. Vault Nanoparticles Packaged with Enzymes as an Efficient Pollutant Biodegradation Technology. ACS Nano 9, 10931–10940 (2015).

4. Wang, Y. & Douglas, T. Tuning properties of biocatalysis using protein cage architectures. J. Mater. Chem. B, 3567–3578 (2023).

5. Wang, Y. & Douglas, T. Protein nanocage architectures for the delivery of therapeutic proteins. Curr. Opin. Colloid Interface Sci. 51, 101395–101395 (2021).

6. McNeale, D. et al. Tunable In Vivo Colocalization of Enzymes within P22 Capsid-Based Nanoreactors. ACS Appl. Mater. Interfaces 15, 17705–17715 (2023).

7. Bonacci, W. et al. Modularity of a carbon-fixing protein organelle. Proc. Natl. Acad. Sci. U. S. A. 109, 478–483 (2012).

8. Frank, S., Lawrence, A.D., Prentice, M.B. & Warren, M.J. Bacterial microcompartments moving into a synthetic biological world. J. Biotechnol. 163, 273–279 (2013).

9. Lau, Y.H., Giessen, T.W., Altenburg, W.J. & Silver, P.A. Prokaryotic nanocompartments form synthetic organelles in a eukaryote. Nat. Commun. 9, 1–7 (2018).

10. Jones, J.A., Andreas, M.P. & Giessen, T.W. Exploring the Extreme Acid Tolerance of a Dynamic Protein Nanocage. Biomacromolecules 24, 1388–1399 (2023).

11. Ebensperger, P. et al. A Dual-Metal-Catalyzed Sequential Cascade Reaction in an Engineered Protein Cage**. Angew. Chemie 135, 1–9 (2023).

12. Tamura, A. et al. Packaging guest proteins into the encapsulin nanocompartment from Rhodococcus erythropolis N771. Biotechnol. Bioeng. 112, 13–20 (2015).

13. Lončar, N., Rozeboom, H.J., Franken, L.E., Stuart, M.C.A. & Fraaije, M.W. Structure of a robust bacterial protein cage and its application as a versatile biocatalytic platform through enzyme encapsulation. Biochem. Biophys. Res. Commun. 529, 548–553 (2020).

14. Sigmund, F. et al. Bacterial encapsulins as orthogonal compartments for mammalian cell engineering. Nat. Commun. 9, 1–14 (2018).

15. Tagit, O., De Ruiter, M.V., Brasch, M., Ma, Y. & Cornelissen, J.J.L.M. Quantum dot encapsulation in virus-like particles with tuneable structural properties and low toxicity. RSC Adv. 7, 38110–38118 (2017).

16. Zhang, J. et al. Cargo loading within ferritin nanocages in preparation for tumor-targeted delivery. Nat. Protoc. 16, 4878–4896 (2021).

17. Stupka, I. et al. Chemically induced protein cage assembly with programmable opening and cargo release. Sci. Adv. 8(2022).

18. Herbert, F.C. et al. Supramolecular Encapsulation of Small-Ultrared Fluorescent Proteins in Virus-Like Nanoparticles for Noninvasive in Vivo Imaging Agents. Bioconjug. Chem. 31, 1529–1536 (2020).

19. Noireaux, V. & Libchaber, A. A vesicle bioreactor as a step toward an artificial cell assembly. Proc. Natl. Acad. Sci. U. S. A. 101, 17669–17674 (2004).

20. Lu, A. & O’Reilly, R.K. Advances in nanoreactor technology using polymeric nanostructures. Curr. Opin. Biotechnol. 24, 639–645 (2013).

21. Jiao, Y., Shang, Y., Li, N. & Ding, B. DNA-based enzymatic systems and their applications. iScience 25, 1–17 (2022).

22. Kalyana Sundaram, S.d., Hossain, M.M., Rezki, M., Ariga, K. & Tsujimura, S. Enzyme Cascade Electrode Reactions with Nanomaterials and Their Applicability towards Biosensor and Biofuel Cells. Biosensors 13(2023).

23. Jutz, G., Van Rijn, P., Santos Miranda, B. & Böker, A. Ferritin: A versatile building block for bionanotechnology. Chem. Rev. 115, 1653–1701 (2015).

24. Su, Y. et al. Virus-like particles nanoreactors: from catalysis towards bio-applications. J. Mater. Chem. B 11, 9084–9098 (2023).

25. Plegaria, J.S. & Kerfeld, C.A. Engineering nanoreactors using bacterial microcompartment architectures. Curr. Opin. Biotechnol. 51, 1–7 (2018).

26. Azuma, Y., Edwardson, T.G.W. & Hilvert, D. Tailoring lumazine synthase assemblies for bionanotechnology. Chem. Soc. Rev. 47, 3543–3557 (2018).

27. Stupka, I. & Heddle, J.G. Artificial protein cages – inspiration, construction, and observation. Vol. 64 66–73 (2020).

28. Jones, J.A. & Giessen, T.W. Advances in encapsulin nanocompartment biology and engineering. Biotechnol. Bioeng. 118, 491–505 (2021).

29. Patterson, D.P. et al. Virus-like particle nanoreactors: Programmed encapsulation of the thermostable CelB glycosidase inside the P22 capsid. Soft Matter 8, 10158–10166 (2012).

30. Uchida, M. et al. Modular Self-Assembly of Protein Cage Lattices for Multistep Catalysis. ACS Nano 12, 942–953 (2018).

31. Jordan, P.C. et al. Self-assembling biomolecular catalysts for hydrogen production. Nat. Chem. 8, 179–185 (2016).

32. González-Davis, O., Chauhan, K., Zapian-Merino, S.J. & Vazquez-Duhalt, R. Bi-enzymatic virus-like bionanoreactors for the transformation of endocrine disruptor compounds. Int. J. Biol. Macromol. 146, 415–421 (2020).

33. Selivanovitch, E., LaFrance, B. & Douglas, T. Molecular exclusion limits for diffusion across a porous capsid. Nat. Commun. 12(2021).

34. Azuma, Y., Bader, D.L.V. & Hilvert, D. Substrate Sorting by a Supercharged Nanoreactor. J. Am. Chem. Soc. 140, 860–863 (2018).

35. Frey, R., Hayashi, T. & Hilvert, D. Enzyme-mediated polymerization inside engineered protein cages. Chem. Commun. 52, 10423–10426 (2016).

36. Lawrence, A.D. et al. Solution Structure of a Bacterial Microcompartment Targeting Peptide and Its Application in the Construction of an Ethanol Bioreactor. ACS Synth. Biol. 3, 454−465-454−465 (2014).

37. Li, T. et al. Reprogramming bacterial protein organelles as a nanoreactor for hydrogen production. Nat. Commun. 11, 1–10 (2020).

38. Jenkins, M.C. & Lutz, S. Encapsulin Nanocontainers as Versatile Scaffolds for the Development of Artificial Metabolons. ACS Synth. Biol. 10, 857–869 (2021).

39. Williams, E.M., Jung, S.M., Coffman, J.L. & Lutz, S. Pore Engineering for Enhanced Mass Transport in Encapsulin Nanocompartments. ACS Synth. Biol. 7, 2514–2517 (2018).

40. Adamson, L.S.R. et al. Pore structure controls stability and molecular flux in engineered protein cages. Sci. Adv. 8, 1–13 (2022).

41. Glasgow, J.E., Asensio, M.A., Jakobson, C.M., Francis, M.B. & Tullman-Ercek, D. Influence of Electrostatics on Small Molecule Flux through a Protein Nanoreactor. ACS Synth. Biol. 4, 1011–1019 (2015).

42. Giessen, T.W. Encapsulins. Annu Rev Biochem (2022).

43. Sutter, M. et al. Structural basis of enzyme encapsulation into a bacterial nanocompartment. Nat Struct Mol Biol 15, 939–47 (2008).

44. McHugh, C.A. et al. A virus capsid-like nanocompartment that stores iron and protects bacteria from oxidative stress. EMBO J 33, 1896–911 (2014).

45. Giessen, T.W. et al. Large protein organelles form a new iron sequestration system with high storage capacity. Elife 8(2019).

46. Eren, E. et al. Structural characterization of the Myxococcus xanthus encapsulin and ferritin-like cargo system gives insight into its iron storage mechanism. Structure 30, 551–563 e4 (2022).

47. Ross, J. et al. Pore dynamics and asymmetric cargo loading in an encapsulin nanocompartment. Sci Adv 8, eabj4461 (2022).

48. Cassidy-Amstutz, C. et al. Identification of a Minimal Peptide Tag for in Vivo and in Vitro Loading of Encapsulin. Biochemistry 55, 3461–8 (2016).

49. Altenburg, W.J., Rollins, N., Silver, P.A. & Giessen, T.W. Exploring targeting peptide-shell interactions in encapsulin nanocompartments. Sci Rep 11, 4951 (2021).

50. Jones, J.A. & Giessen, T.W. Advances in encapsulin nanocompartment biology and engineering. Biotechnol Bioeng 118, 491–505 (2021).

51. Gabashvili, A.N. et al. Encapsulins-Bacterial Protein Nanocompartments: Structure, Properties, and Application. Biomolecules 10(2020).

52. Sigmund, F. et al. Bacterial encapsulins as orthogonal compartments for mammalian cell engineering. Nat Commun 9, 1990 (2018).

53. Smart, O.S., Neduvelil, J.G., Wang, X., Wallace, B.A. & Sansom, M.S.P. HOLE: A program for the analysis of the pore dimensions of ion channel structural models. J. Mol. Graph. 14, 354–360 (1996).

54. Szyszka, T.N., et al. Point mutation in a virus-like capsid drives symmetry reduction to form tetrahedral cages. bioRxiv (2024).

55. Edwardson, T.G.W., Mori, T. & Hilvert, D. Rational Engineering of a Designed Protein Cage for siRNA Delivery. J Am Chem Soc 140, 10439–10442 (2018).

56. Lopez-Otin, C. & Bond, J.S. Proteases: multifunctional enzymes in life and disease. J Biol Chem 283, 30433–7 (2008).

57. Yu, W.H., Huang, P.T., Lou, K.L., Yu, S.S. & Lin, C. A smallest 6 kda metalloprotease, mini-matrilysin, in living world: a revolutionary conserved zinc-dependent proteolytic domain-helix-loop-helix catalytic zinc binding domain (ZBD). J Biomed Sci 19, 54 (2012).

58. Sharma, J. & Douglas, T. Tuning the catalytic properties of P22 nanoreactors through compositional control. Nanoscale 12, 336–346 (2020).

59. Zou, Y. et al. Crystal structures of phosphite dehydrogenase provide insights into nicotinamide cofactor regeneration. Biochemistry 51, 4263–70 (2012).

60. Smart, O.S., Goodfellow, J.M. & Wallace, B.A. The pore dimensions of gramicidin A. Biophys J 65, 2455–60 (1993).

61. Suloway, C. et al. Automated molecular microscopy: the new Leginon system. J Struct Biol 151, 41–60 (2005).

62. Punjani, A., Rubinstein, J.L., Fleet, D.J. & Brubaker, M.A. cryoSPARC: algorithms for rapid unsupervised cryo-EM structure determination. Nat Methods 14, 290–296 (2017).

63. Pettersen, E.F. et al. UCSF ChimeraX: Structure visualization for researchers, educators, and developers. Protein Science 30, 70–82 (2021).

64. Emsley, P., Lohkamp, B., Scott, W.G. & Cowtan, K. Features and development of Coot. Acta Crystallographica Section D: Biological Crystallography 66, 486–501 (2010).

