## Supplementary Information for "Pore engineering as a general strategy to improve protein-based enzyme nanoreactor performance"

### Table of Contents

Supplementary Fig. 1. SDS-PAGE gels of all proteins used in this study. 2
Supplementary Fig. 2. Additional negative-stain TEM micrographs 3
Supplementary Fig. 3. Cryo-EM processing of Mx_pmut T=1 shells 4
Supplementary Fig. 4. Cryo-EM processing of Mx_pmut T=3 and T=4 shells 5
Supplementary Fig. 5. Local resolution analysis of Mx_pmut shells 6
Supplementary Fig. 6. Dynamic light scattering analysis and thermal ramps 7
Supplementary Fig. 7. Trypsin and bacterial cell lysate incubations 10
Supplementary Fig. 8. Activity analysis of co-encapsulated PTDH 12
Supplementary Fig. 9. Activity analysis of co-encapsulated AdhD 13
Supplementary Table 1. Sequences of constructs used in this study 14
Supplementary Table 2. DNA primers used in this study 19
Supplementary Table 3. Cryo-EM data collection and refinement statistics 20

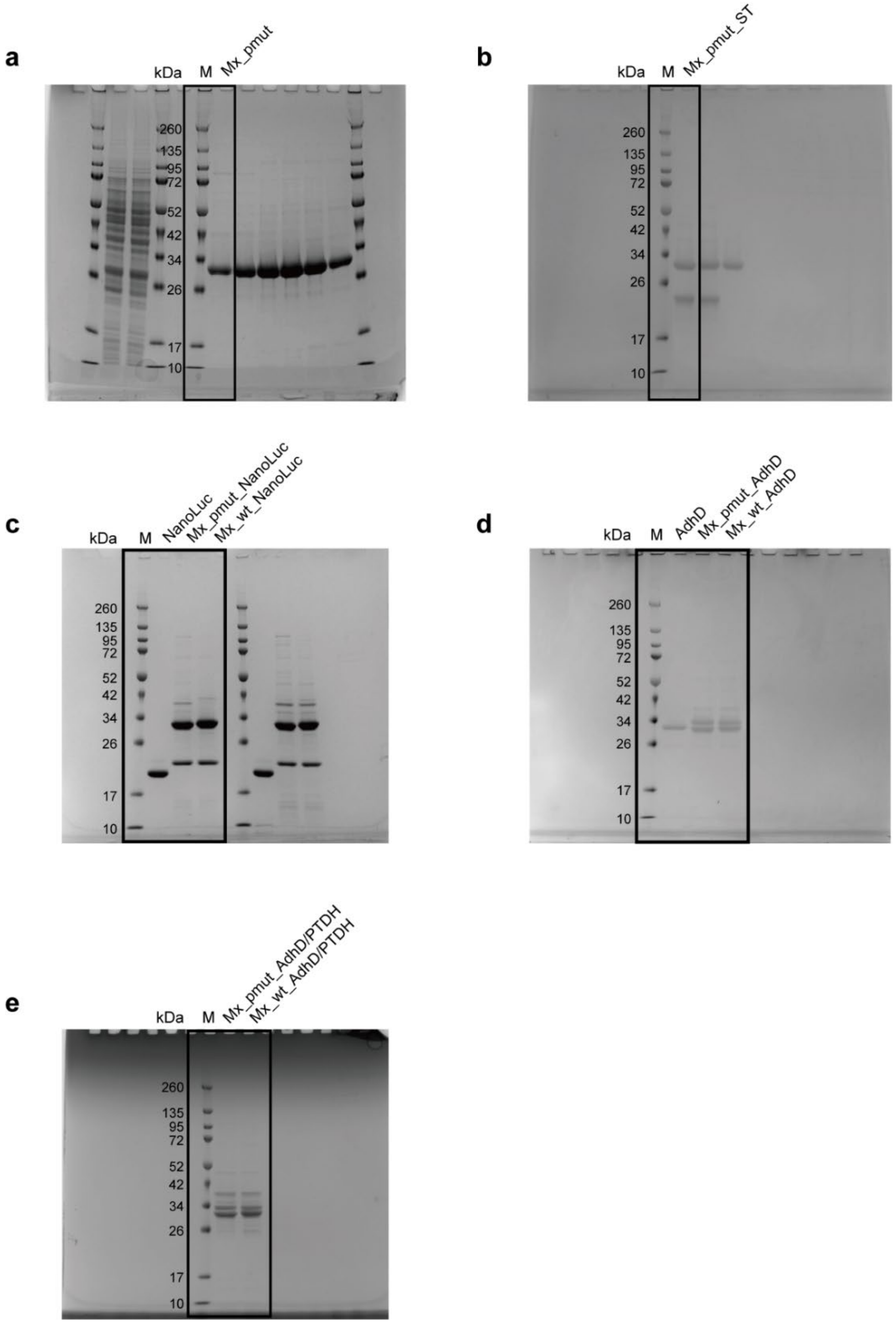

**Supplementary Fig. 1. Full uncropped SDS-PAGE gels of all purified proteins used in this study.** **a**, Mx_pmut. **b**, SNAP-tag (ST)-loaded Mx_pmut. **c**, proteins and nanoreactors for NanoLuc experiments. **d**, proteins and nanoreactors for AdhD experiments. **e**, proteins and nanoreactors used for cofactor recycling experiments.

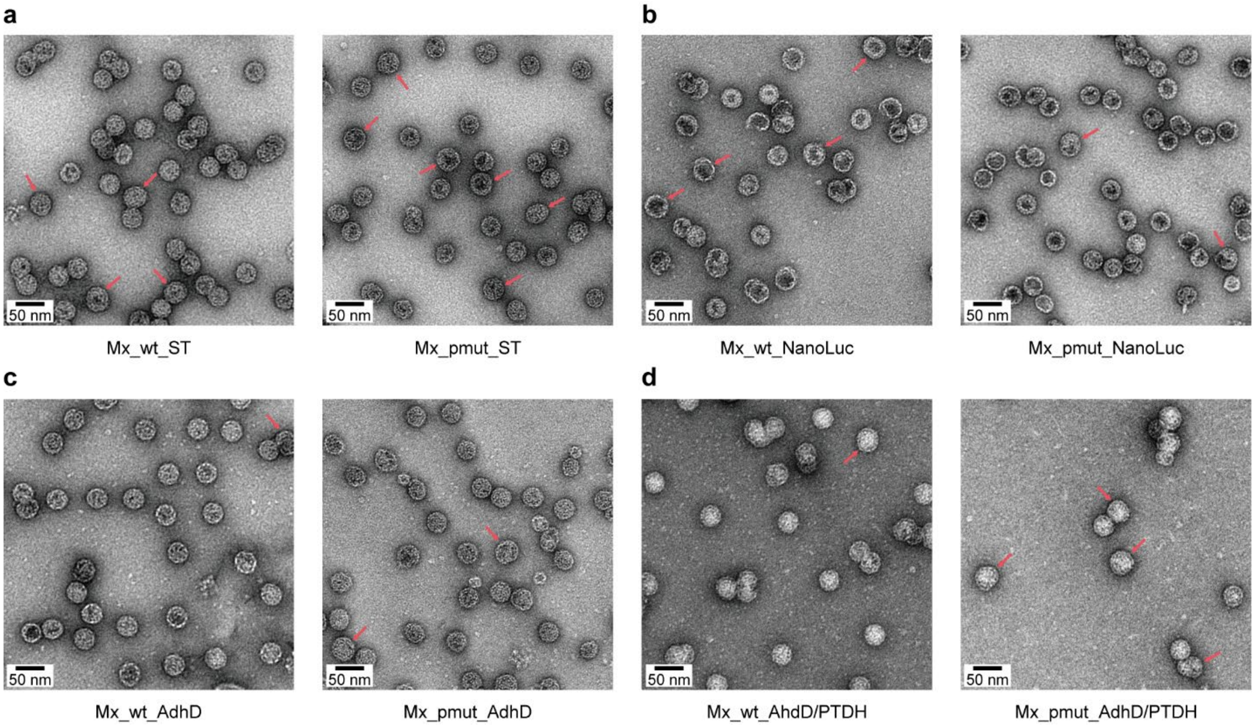

**Supplementary Fig. 2. Negative-stain TEM micrographs of Mx_pmut- and Mx_wt-based-nanoreactors. a**, SNAP-tag (ST) cargo. **b**, NanoLuc cargo. **c**, AdhD cargo. **d**, AdhD/PTDH cargo. Likely T=4 shells are highlighted with red arrows.

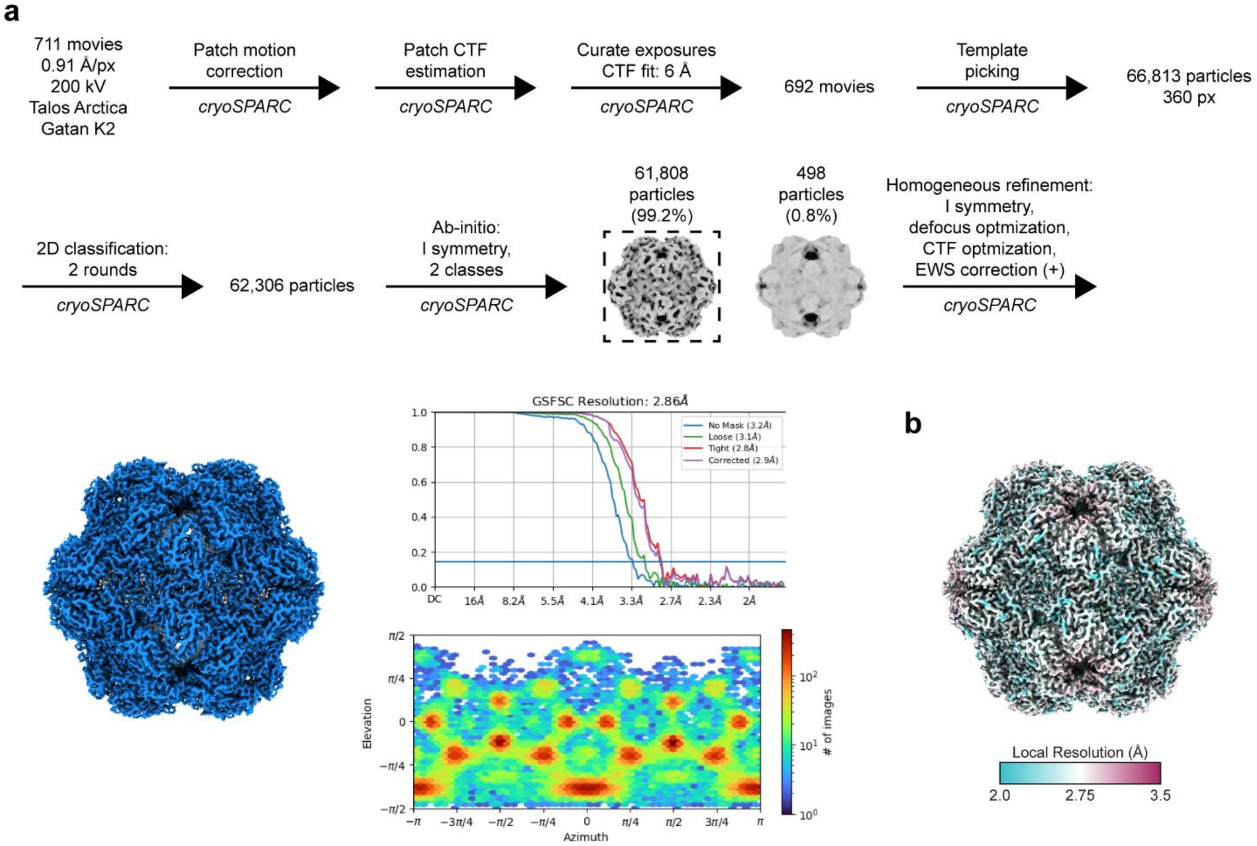

**Supplementary Fig 3. Cryo-EM processing and analysis of Mx_pmut T=1 shells. a**, Cryo-EM data processing workflow for the Mx_pmut T=1 shell without cargo. GSFSC: gold-standard Fourier shell correlation. Angular distribution of particles shown below the GSFSC plot. **b**, Local resolution analysis of the Mx_pmut T=1 cryo-EM density.

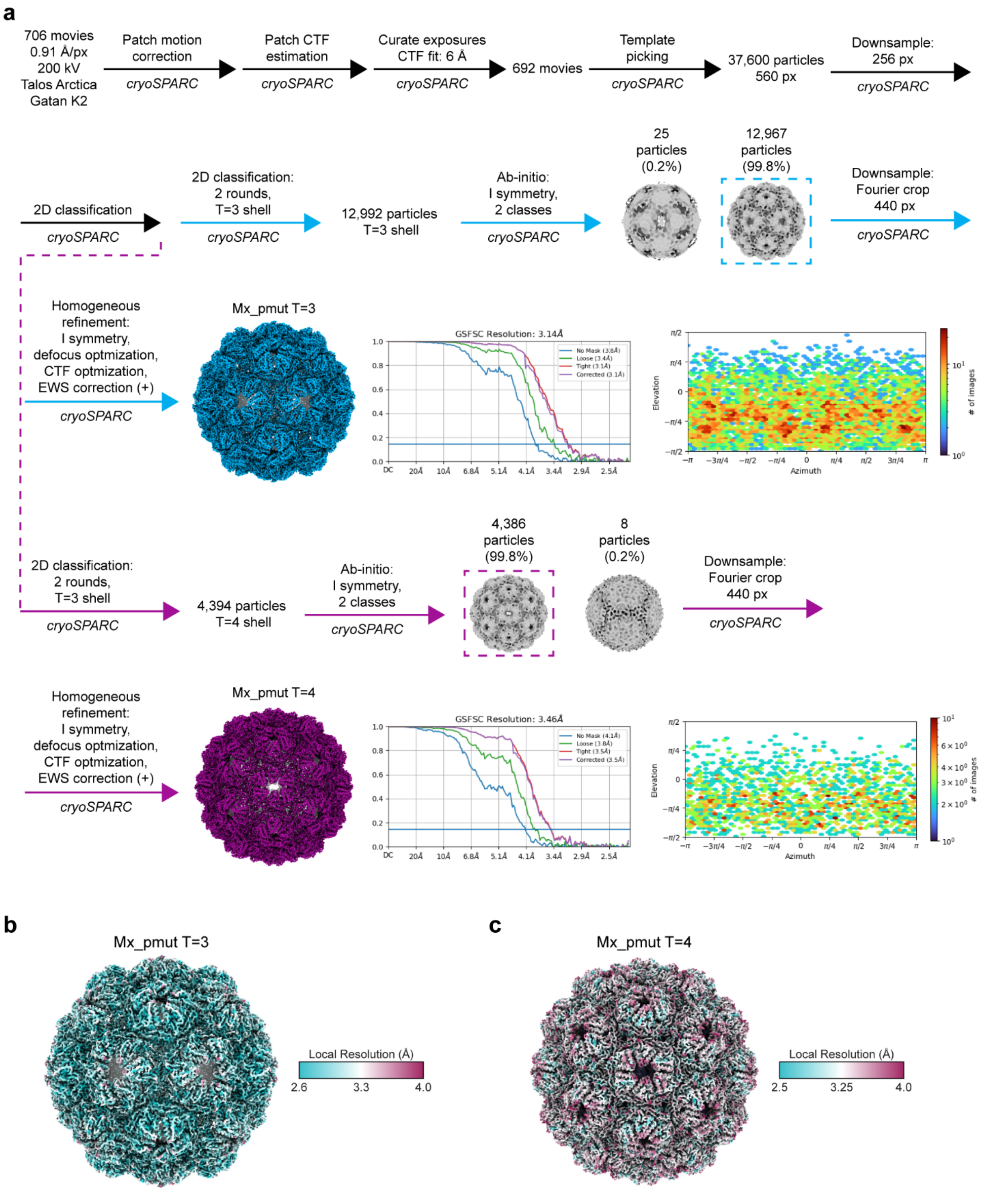

**Supplementary Fig. 4. Cryo-EM processing and analysis of Mx_pmut T=3 and T=4 shells.** **a**, Cryo-EM data processing workflow for Mx_pmut T=3 (cyan) and Mx_pmut T=4 (magenta). Angular distribution of particles shown below the GSFSC plots. **b**, Local resolution analysis of the Mx_pmut T=3 cryo-EM density. **c**, Local resolution analysis of the T=4 cryo-EM density.

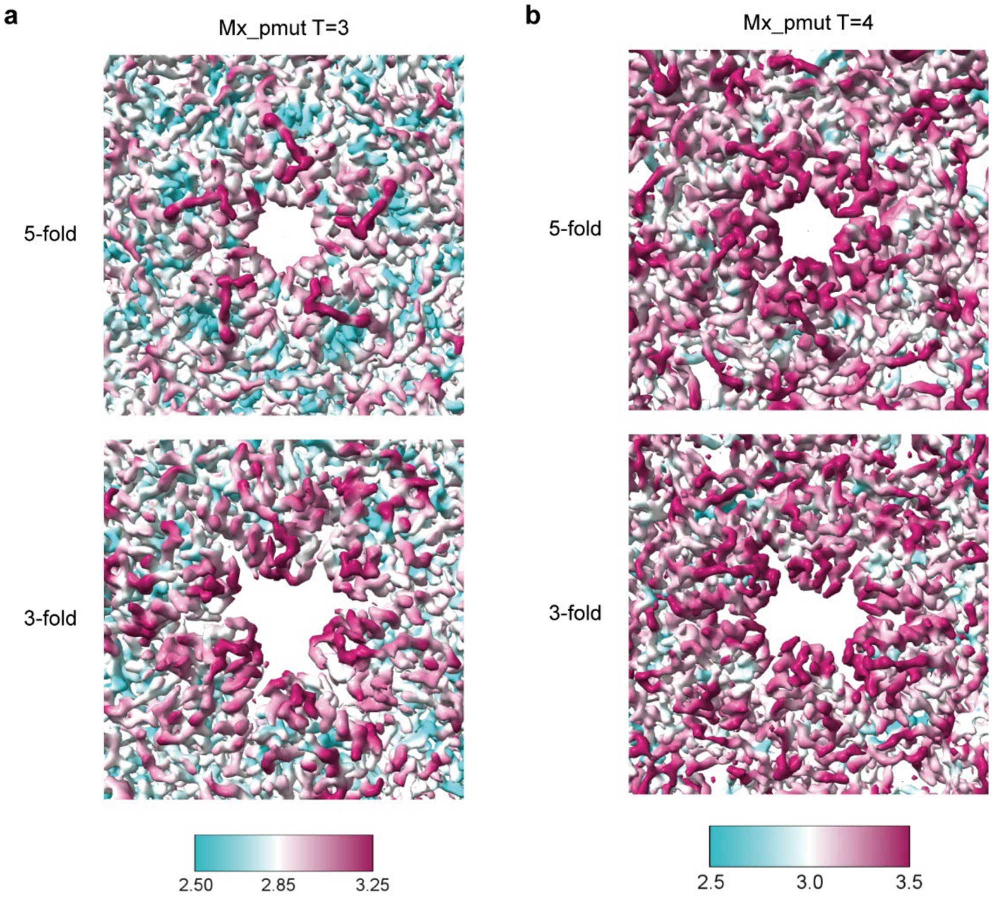

**Supplementary Fig. 5. Local resolution analysis of Mx_pmut T=3 and T=4 pores. a**, Mx_pmut T=3 5- and 3-fold pores. Local resolution scale shown below in Å. **b**, Mx_pmut T=4 5- and 3-fold pores. Local resolution scale shown below in Å.

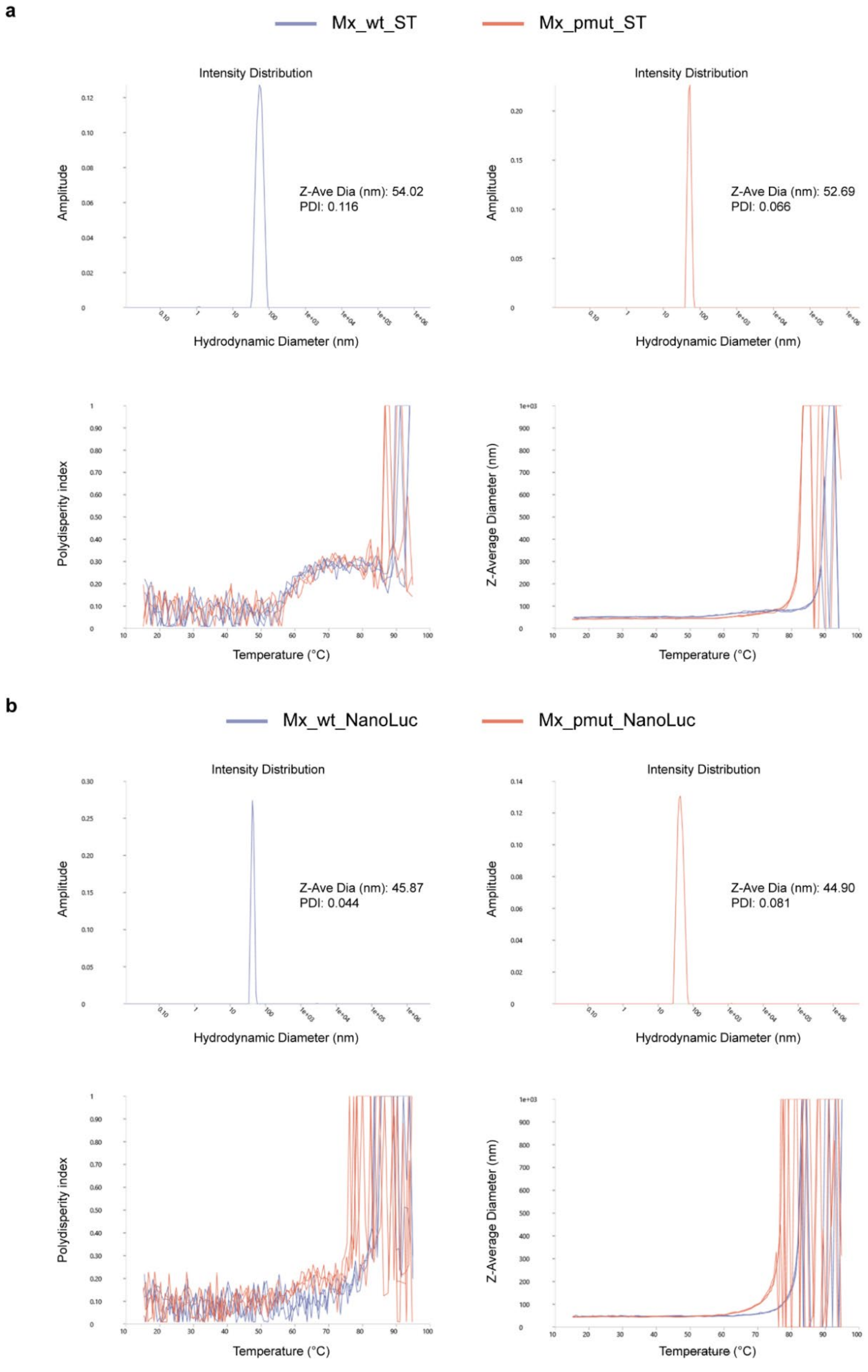
a
Mx_wt_ST
Mx_pmut_ST
Intensity Distribution
Amplitude
Z-Ave Dia (nm): 54.02
PDI: 0.116
Hydrodynamic Diameter (nm)
Intensity Distribution
Amplitude
Z-Ave Dia (nm): 52.69
PDI: 0.066
Hydrodynamic Diameter (nm)
Polydisperity index
Temperature (°C)
Z-Average Diameter (nm)
Temperature (°C)
b
Mx_wt_NanoLuc
Mx_pmut_NanoLuc
Intensity Distribution
Amplitude
Z-Ave Dia (nm): 45.87
PDI: 0.044
Hydrodynamic Diameter (nm)
Intensity Distribution
Amplitude
Z-Ave Dia (nm): 44.90
PDI: 0.081
Hydrodynamic Diameter (nm)
Polydisperity index
Temperature (°C)
Z-Average Diameter (nm)
Temperature (°C)

c

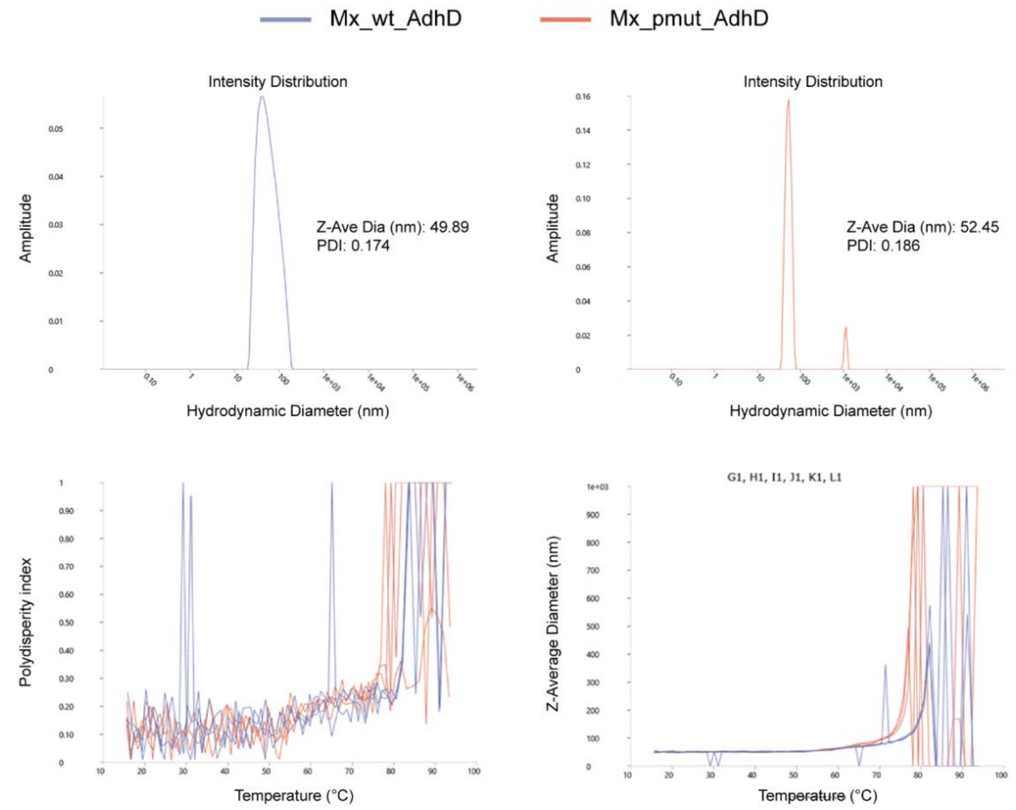

Mx_wt_AdhD
Mx_pmut_AdhD
Intensity Distribution
Amplitude
Z-Ave Dia (nm): 49.89
PDI: 0.174
Hydrodynamic Diameter (nm)
Intensity Distribution
Amplitude
Z-Ave Dia (nm): 52.45
PDI: 0.186
Hydrodynamic Diameter (nm)
Polydisperity index
Temperature (°C)
G1, H1, I1, J1, K1, L1
Z-Average Diameter (nm)
Temperature (°C)

d

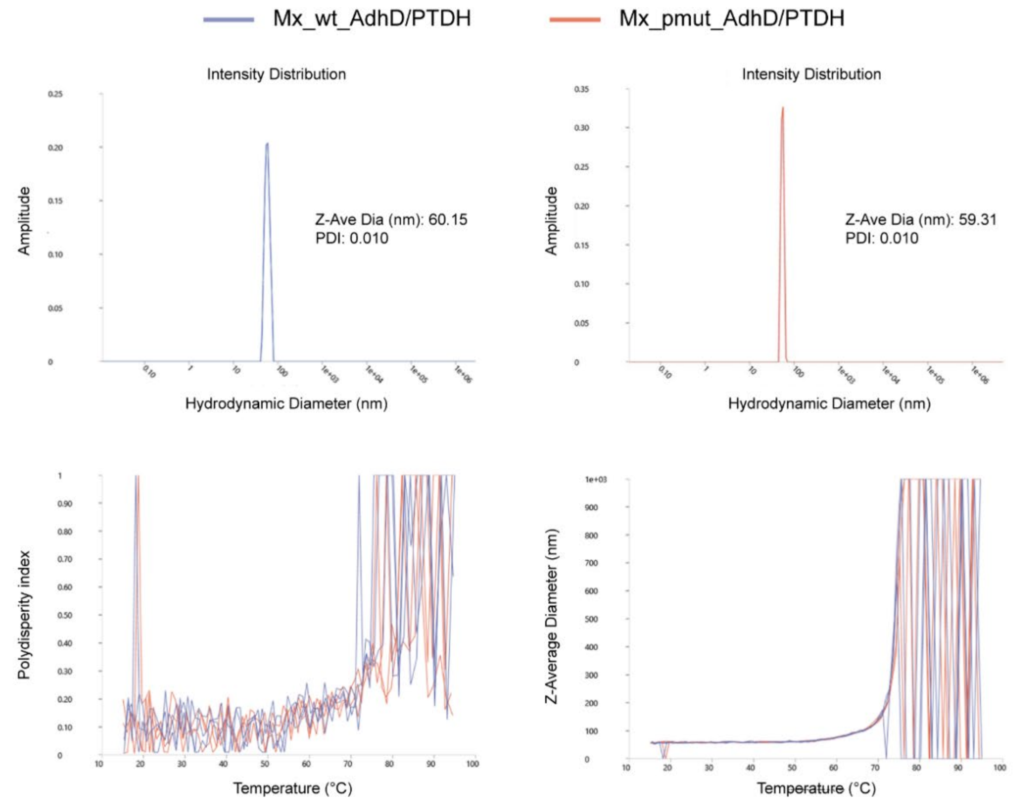

Mx_wt_AdhD/PTDH
Mx_pmut_AdhD/PTDH
Intensity Distribution
Amplitude
Z-Ave Dia (nm): 60.15
PDI: 0.010
Hydrodynamic Diameter (nm)
Intensity Distribution
Amplitude
Z-Ave Dia (nm): 59.31
PDI: 0.010
Hydrodynamic Diameter (nm)
Polydisperity index
Temperature (°C)
Z-Average Diameter (nm)
Temperature (°C)

**Supplementary Fig. 6. Dynamic light scattering (DLS) analyses coupled with thermal ramps for Mx_wt- and Mx_pmut-based nanoreactors.** **a**, SNAP-tag (ST) cargo. **b**, NanoLuc cargo. **c**, AdhD cargo. **d**, AdhD/PTDH cargo. Z-average diameters with polydispersity index (PDI) values for each nanoreactor system are shown (top left and right). Temperature was increased from 15°C to 95°C at 1°C/min while measuring PDI (bottom left) and Z-average diameter (bottom right). There was no meaningful thermal stability difference between Mx_wt- and Mx_pmut-based nanoreactors.

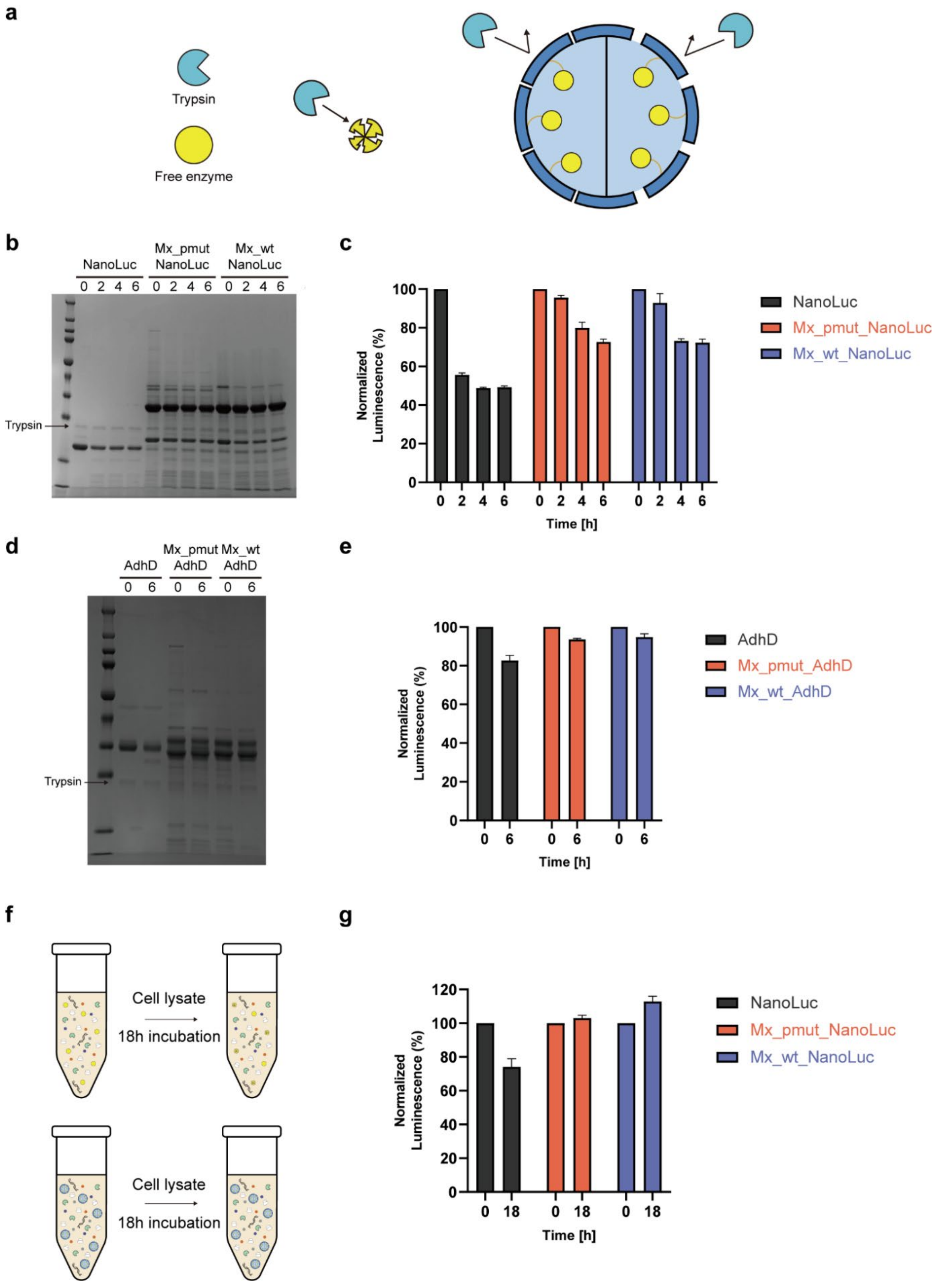

a
Trypsin
Free enzyme
b
NanoLuc
Mx_pmut NanoLuc
Mx_wt NanoLuc
0 2 4 6 0 2 4 6 0 2 4 6
Trypsin
c
Normalized Luminescence (%)
Time [h]
NanoLuc
Mx_pmut_NanoLuc
Mx_wt_NanoLuc
d
AdhD
Mx_pmut AdhD
Mx_wt AdhD
0 6 0 6 0 6
Trypsin
e
Normalized Luminescence (%)
Time [h]
AdhD
Mx_pmut_AdhD
Mx_wt_AdhD
f
Cell lysate
18h incubation
Cell lysate
18h incubation
g
Normalized Luminescence (%)
Time [h]
NanoLuc
Mx_pmut_NanoLuc
Mx_wt_NanoLuc

**Supplementary Fig. 7. Trypsin and bacterial cell lysate incubations. a**, Schematic illustrating the susceptibility of free enzymes to trypsin and the protective effect of Mx_pmut and Mx_wt shells. **b**, SDS-PAGE analysis showing the proteolytic resistance of Mx_wt- and Mx_pmut-encapsulated NanoLuc compared to free NanoLuc upon trypsin exposure. Band intensity for free NanoLuc markedly reduced following a 2 h incubation period while Mx_wt- and Mx_pmut-encapsulated NanoLuc exhibited little loss of band intensity over the examined time course. **c**, Enzymatic activity of trypsin-incubated free and Mx_wt- and Mx_pmut-encapsulated NanoLuc at given time points normalized by the activity before incubation. The activity of encapsulated NanoLuc was substantially more retained than that of free NanoLuc. **d**, SDS-PAGE analysis showing the proteolytic resistance of Mx_wt- and Mx_pmut-encapsulated AdhD compared to free AdhD upon trypsin exposure. Compared to NanoLuc, free AdhD was more resistant to trypsin exposure. **e**, Normalized enzymatic activity of trypsin-incubated free and Mx_wt- and Mx_pmut-encapsulated AdhD after 6 h of incubation. **f**, Schematic illustrating the susceptibility of free enzymes to harsh bacterial lysate conditions containing various proteases and reactive molecules and the protective effect of enzyme encapsulation. **g**, Normalized enzymatic activity of bacterial lysate-incubated free and Mx_wt- and Mx_pmut-encapsulated NanoLuc after 18 h of incubation. All error bars represent standard errors from three independent experiments.

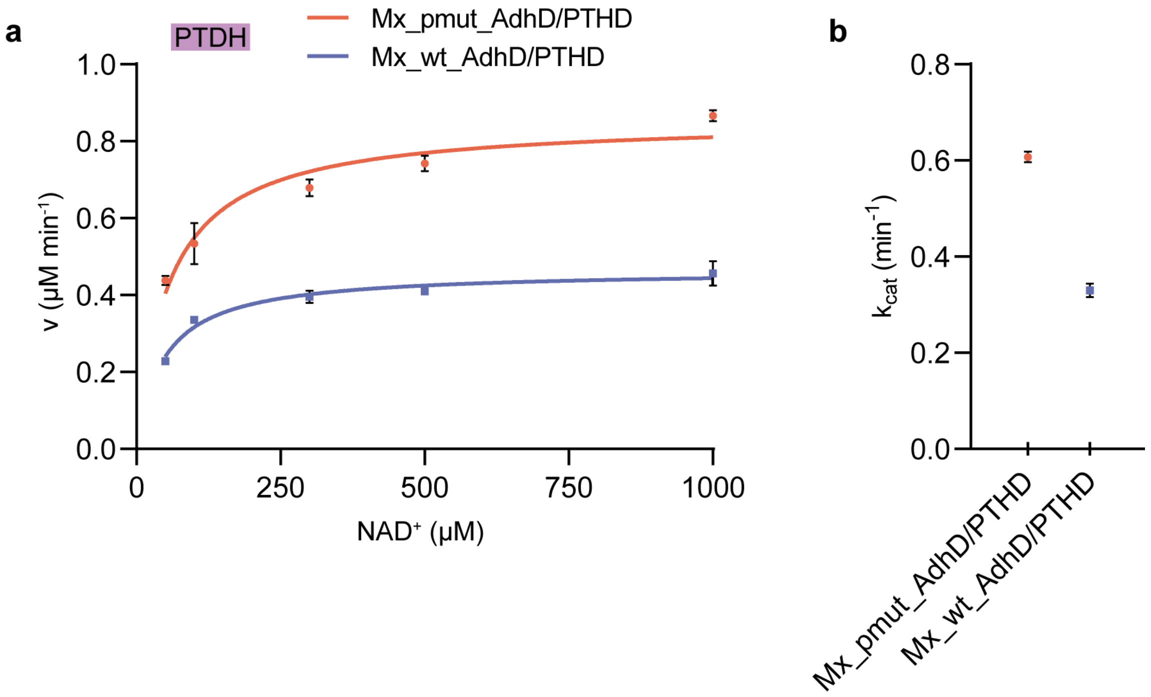

**Supplementary Fig. 8. Activity analysis of co-encapsulated PTDH.** **a**, Saturation kinetics analysis of PTDH in Mx_pmut- and Mx_wt-based AdhD/PTDH co-encapsulation nanoreactors. **b**, Comparison of turnover numbers ($k_{cat}$) for PTDH in Mx_pmut- and Mx_wt-based AdhD/PTDH co-encapsulation nanoreactors. All error bars represent standard error of three independent experiments.

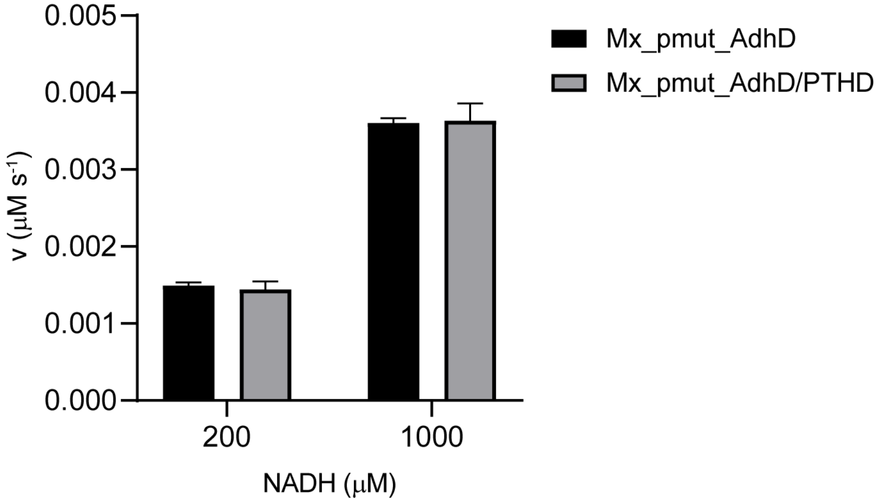

**Supplementary Fig. 9. Activity analysis of co-encapsulated AdhD.** AdhD activity assay examining the impact of co-encapsulation on AdhD activity compared to AdhD-only nanoreactors. For both nanoreactors, final concentrations of encapsulated AdhD in the reaction mixture was 15 nM and the AdhD activity was measured with 200 μM and 1000 μM NADH. There was no difference in AdhD activity demonstrating that AdhD activity was not negatively affected by co-encapsulation.

**Supplementary Table 1. DNA and amino acid sequences of proteins used in this study.**

| Construct name | DNA sequence | Protein sequence |
|---|---|---|
| Mx_wt | ATGCCAGATTTTTTGGGTCACGCAGAAAAC<br>CCACTCCGGGAGGAAGAGTGGGCTCGCCT<br>TAACGAGACAGTTATTCAGGTTGCCCGTCG<br>TTCGCTTGTGGGGCGTCGGATTCTTGATAT<br>TTATGGCCCTTTGGGCGCAGGCGTCCAAAC<br>AGTACCATATGACGAATTTCAGGGTGTGAG<br>CCCAGGCGCAGTAGACATCGTCGGGGAAC<br>AAGAAACTGCTATGGTCTTCACCGACGCCC<br>GGAAGTTCAAAACTATCCCTATCATTTACAA<br>AGACTTCCTCCTTCATTGGCGTGACATCGA<br>AGCTGCGCGCACGCATAATATGCCTCTTGA<br>TGTAAGTGCGGCCGCCGGTGCAGCTGCCC<br>TTTGCGCCCAGCAAGAGGATGAGCTTATCT<br>TCTATGGCGACGCTCGGCTCGGGTATGAA<br>GGCCTTATGACGGCGAACGGTCGGCTGAC<br>TGTTCCATTAGGTGACTGGACTTCCCCGGG<br>TGGCGGCTTCCAGGCCATTGTCGAAGCCA<br>CTCGTAAGTTAAACGAACAAGGCCACTTTG<br>GTCCATACGCTGTTGTGCTGTCACCTCGCT<br>TATATTCCCAGTTACATCGGATTTACGAAAA<br>AACAGGGGTCTTAGAGATCGAGACAATTCG<br>CCAGCTCGCCTCAGATGGTGTTTATCAGTC<br>GAATCGGTTACGGGGTGAGAGTGGCGTCG<br>TGGTCTCTACAGGCCGTGAAAACATGGATT<br>TAGCGGTGAGTATGGATATGGTTGCAGCCT<br>ACTTAGGGGCATCCCGGATGAATCACCCTT<br>TTCGCGTACTGGAAGCCCTCCTTTTGCGCA<br>TCAAGCATCCTGACGCGATCTGTACGTTAG<br>AAGGTGCTGGTGCGACTGAGCGGCGC | MPDFLGHAENPLREEEWARLNETVIQVAR<br>RSLVGRRILDIYGPLGAGVQTVPYDEFQGV<br>SPGAVDIVGEQETAMVFTDARKFKTIPIIYK<br>DFLLHWRDIEAARTHNMPLDVSAAAGAAAL<br>CAQQEDELIFYGDARLGYEGLMTANGRLT<br>VPLGDWTSPGGGFQAIVEATRKLNEQGHF<br>GPYAVVLSPRLYSQLHRIYEKTGVLEIETIR<br>QLASDGVYQSNRLRGESGVVVSTGRENM<br>DLAVSMDMVAAYLGASRMNHPFRVLEALL<br>LRIKHPDAICTLEGAGATERR |
| Mx_pmut | ATGCCAGATTTTTTGGGTCACGCAGAAAAC<br>CCACTCCGGGAGGAAGAGTGGGCTCGCCT<br>TAACGAGACAGTTATTCAGGTTGCCCGTCG<br>TTCGCTTGTGGGGCGTCGGATTCTTGATAT<br>TTATGGCCCTTTGGGCGCAGGCGTCCAAAC<br>AGTACCATATGACGAATTTCAGGGTGTGAG<br>CCCAGGCGCAGTAGACATCGTCGGGGAAC<br>AAGAAACTGCTATGGTCTTCACCGACGCCC<br>GGAAGTTCAAAACTATCCCTATCATTTACAA<br>AGACTTCCTCCTTCATTGGCGTGACATCGA<br>AGCTGCGCGCACGCATAATATGCCTCTTGA<br>TGTAAGTGCGGCCGCCGGTGCAGCTGCCC<br>TTTGCGCCCAGCAAGAGGATGAGCTTATCT<br>TCTATGGCGACGCTCGGCTCGGGTATGAA<br>GGCCTTATGACGGCGAACGGTCGGCTGAC<br>TGTTCCATTAGGTGACTGGACTTCCCCGGG<br>TGGCGGCTTCCAGGCCATTGTCGAAGCCA<br>CTCGTAAGTTAAACGAACAAGGCCACTTTG<br>GTCCATACGCTGTTGTGCTGTCACCTCGCT<br>TATATTCCCAGTTACATCGGGGTGGCGAGA<br>TCGAGACAATTCGCCAGCTCGCCTCAGATG<br>GTGTTTATCAGTCGAATCGGTTACGGGGTG<br>AGAGTGGCGTCGTGGTCTCTACAGGCCGT<br>GAAAACATGGATTTAGCGGTGAGTATGGAT<br>ATGGTTGCAGCCTACTTAGGGGCATCCCG<br>GATGAATCACCCTTTTCGCGTACTGGAAGC<br>CCTCCTTTTGCGCATCAAGCATCCTGACGC<br>GATCTGTACGTTAGAAGGTGCTGGTGCGAC<br>TGAGCGGCGC | MPDFLGHAENPLREEEWARLNETVIQVAR<br>RSLVGRRILDIYGPLGAGVQTVPYDEFQGV<br>SPGAVDIVGEQETAMVFTDARKFKTIPIIYK<br>DFLLHWRDIEAARTHNMPLDVSAAAGAAAL<br>CAQQEDELIFYGDARLGYEGLMTANGRLT<br>VPLGDWTSPGGGFQAIVEATRKLNEQGHF<br>GPYAVVLSPRLYSQLHRGGEIETIRQLASD<br>GVYQSNRLRGESGVVVSTGRENMDLAVS<br>MDMVAAYLGASRMNHPFRVLEALLLRIKHP<br>DAICTLEGAGATERR |
| SNAP-tag_His | ATGGGCCCAGGGAGTGACAAGGATTGCGA<br>GATGAAGCGGACCACTTTGGACTCACCGCT<br>CGGGAAGCTTGAATTGTCCGGTTGCGAGC<br>AAGGGTTACACGAGATTATCTTTTTAGGTAA<br>GGGGACGAGTGCTGCGGACGCAGTAGAAG | MGPGSDKDCEMKRTTLDSPLGKLELSGCE<br>QGLHEIIFLGKGTSAADAVEVPAPAAVLGG<br>PEPLMQATAWLNAYFHQPEAIEEFPVPALH<br>HPVFQQESFTRQVLWKLLKVVKFGEVISYS<br>HLAALAGNPAATAAVKTALSGNPVPILIPCH |

| | | |
|---|---|---|
| | TTCCTGCTCCTGCCGCGGTGTTGGGCGGT CCTGAACCGCTCATGCAAGCTACGGCGTG GCTCAACGCCTATTTTCACCAGCCGGAAGC TATCGAAGAGTTTCCGGTACCTGCATTGCA CCATCCTGTGTTCCAACAAGAGTCTTTCAC ACGCCAGGTCCTGTGGAAGTTATTAAAGGT TGTCAAGTTTGGCGAGGTCATCAGCTATAG TCACTTGGCGGCCCTCGCTGGTAATCCTGC TGCCACAGCCGCGGTAAAAACTGCACTCTC CGGTAACCCTGTACCGATTTTAATCCCGTG CCACCGTGTGGTTCAGGGGGATCTGGATG TCGGGGGCTACGAGGGGGGGTTAGCGGTT AAGGAATGGTTGCTCGCTCATGAAGGCCAT CGTTTGGGGAAACGGGGCGGGGGTTCAGG CGGGGGCAGTCATCACCACCATCACCAT | RVVQGDLDVGGYEGGLAVKEWLLAHEGH RLGKRGGGSGGGSHHHHHH |
| SNAP-tag_MxTP | ATGGGTCCAGGTAGTGATAAAGATTGCGAA ATGAAACGCACTACCTTAGATTCCCCGCTC GGTAAGTTGGAGTTGAGCGGGTGCGAACA AGGCTTGCATGAAATCATTTTTCTGGGTAA GGGCACGTCAGCAGCAGATGCAGTAGAAG TGCCAGCCCCGGCGGCTGTTCTGGGGGGC CCAGAGCCGCTGATGCAAGCTACCGCATG GCTCAATGCCTATTTTCACCAGCCTGAGGC TATCGAGGAGTTTCCTGTGCCTGCGCTTCA CCATCCGGTCTTCCAGCAGGAGTCGTTTAC TCGGCAGGTACTGTGGAAGCTGTTGAAGGT TGTCAAGTTTGGGGAAGTAATCTCATATAGT CACTTAGCCGCGTTAGCCGGTAATCCTGCG GCTACAGCCGCGGTGAAGACTGCTTTAAGC GGGAACCCAGTTCCTATCCTTATCCCATGC CACCGCGTGGTACAAGGCGACCTCGACGT GGGCGGGTATGAGGGTGGGTTGGCGGTCA AGGAATGGTTGCTGGCCCATGAAGGGCAT CGGTTGGGGAAACGTGGTGGGGGTAGTGG CGGGGGTAGCCCAGAAAAGCGCTTGACAG TAGGGTCTCTCCGGCGT | MGPGSDKDCEMKRTTLDSPLGKLELSGCE QGLHEIIFLGKGTSAADAVEVPAPAAVLGG PEPLMQATAWLNAYFHQPEAIEEFPVPALH HPVFQQESFTRQVLWKLLKVVKFGEVISYS HLAALAGNPAATAAVKTALSGNPVPILIPCH RVVQGDLDVGGYEGGLAVKEWLLAHEGH RLGKRGGGSGGGSPEKRLTVGSLRR |
| NanoLuc_His | ATGGTTTTCACCTTAGAAGATTTCGTGGGC GACTGGCGCCAGACTGCAGGTTACAATCTC GACCAAGTACTCGAGCAGGGTGGTGTCTC GAGCTTATTTCAAAATCTGGGCGTGAGTGT GACTCCGATTCAGCGTATCGTACTGTCCGG TGAGAACGGGCTGAAGATCGACATCCACGT GATCATCCCGTACGAAGGGCTCTCGGGCG ATCAGATGGGCCAAATTGAGAAAATCTTTAA GGTGGTTTACCCGGTTGATGACCATCATTT TAAAGTTATCCTGCATTACGGCACGCTTGT CATTGACGGCGTTACACCAAACATGATCGA CTACTTTGGGCGCCCGTACGAAGGGATCG CTGTGTTTGACGGGAAAAAAATCACCGTAA CGGGTACTCTCTGGAATGGTAATAAGATTA TCGACGAACGTTTAATCAATCCAGACGGCT CTCTCTTGTTCCGCGTCACTATCAATGGCG TAACAGGTTGGCGTTTATGTGAGCGTATCC TCGCCGGTGGTGGGAGCGGGGGCGGTTC ACATCACCACCATCACCAT | MVFTLEDFVGDWRQTAGYNLDQVLEQGG VSSLFQNLGVSVTPIQRIVLSGENGLKIDIHV IIPYEGLSGDQMGQIEKIFKVVYPVDDHHFK VILHYGTLVIDGVTPNMIDYFGRPYEGIAVF DGKKITVTGTLWNGNKIIDERLINPDGSLLF RVTINGVTGWRLCERILAGGGSGGGSHHH HHH |
| NanoLuc_MxTP | ATGGTTTTCACCTTAGAAGATTTCGTGGGC GACTGGCGCCAGACTGCAGGTTACAATCTC GACCAAGTACTCGAGCAGGGTGGTGTCTC GAGCTTATTTCAAAATCTGGGCGTGAGTGT GACTCCGATTCAGCGTATCGTACTGTCCGG TGAGAACGGGCTGAAGATCGACATCCACGT GATCATCCCGTACGAAGGGCTCTCGGGCG ATCAGATGGGCCAAATTGAGAAAATCTTTAA GGTGGTTTACCCGGTTGATGACCATCATTT TAAAGTTATCCTGCATTACGGCACGCTTGT CATTGACGGCGTTACACCAAACATGATCGA CTACTTTGGGCGCCCGTACGAAGGGATCG | MVFTLEDFVGDWRQTAGYNLDQVLEQGG VSSLFQNLGVSVTPIQRIVLSGENGLKIDIHV IIPYEGLSGDQMGQIEKIFKVVYPVDDHHFK VILHYGTLVIDGVTPNMIDYFGRPYEGIAVF DGKKITVTGTLWNGNKIIDERLINPDGSLLF RVTINGVTGWRLCERILAGGGSGGGSPEK RLTVGSLRR |

| | | |
|---|---|---|
| | CTGTGTTTGACGGGAAAAAAATCACCGTAA<br>CGGGTACTCTCTGGAATGGTAATAAGATTA<br>TCGACGAACGTTTAATCAATCCAGACGGCT<br>CTCTCTTGTTCCGCGTCACTATCAATGGCG<br>TAACAGGTTGGCGTTTATGTGAGCGTATCC<br>TCGCCGGTGGTGGGAGCGGGGGCGGTTC<br>ACCTGAGAAACGCTTGACTGTTGGGTCTCT<br>CCGGCGT | |
| AdhD_His | ATGAAGCGCGTTAACGCATTTAACGACCTG<br>AAACGTATTGGGGATGATAAGGTTACTGCA<br>ATTGGTATGGGGACATGGGGCATCGGGGG<br>CCGTGAAACTCCTGACTATTCCCGCGACAA<br>GGAGTCCATCGAAGCAATCCGGTATGGCTT<br>GGAGCTCGGCATGAACCTTATTGACACGGC<br>TGAATTTTATGGGGCGGGCCACGCAGAAG<br>AGATTGTTGGTGAGGCTATCAAGGAATTTG<br>AACGTGAGGACATTTTTATCGTGTCGAAGG<br>TATGGCCGACACATTTTGGGTACGAAGAAG<br>CAAAGAAAGCGGCCCGGGCCTCGGCCAAG<br>CGCTTGGGTACTTATATTGATTTGTACTTAT<br>TGCACTGGCCTGTTGACGATTTCAAAAAAA<br>TTGAAGAGACCTTGCATGCCCTTGAAGACC<br>TCGTCGACGAGGGGGTCATCCGTTACATC<br>GGCGTTTCAAACTTCAACTTAGAACTTCTCC<br>AACGGTCCCAGGAGGTGATGCGCAAATAC<br>GAGATCGTAGCGAACCAAGTGAAGTATTCT<br>GTAAAGGACCGGTGGCCAGAAACCACGGG<br>TCTCCTGGATTACATGAAGCGGGAGGGCAT<br>CGCTTTAATGGCTTACACGCCACTTGAGAA<br>AGGTACTTTGGCGCGGAATGAATGTCTTGC<br>GAAGATCGGGGAGAAATACGGTAAAACCG<br>CGGCACAGGTAGCCCTCAACTATCTGATTT<br>GGGAAGAAAATGTGGTGGCGATCCCTAAG<br>GCATCTAATAAGGAACACTTGAAAGAGAAT<br>TTCGGGGCCATGGGTTGGCGCCTTAGCGA<br>AGAAGACCGCGAGATGGCTCGTCGCTGTG<br>TTGGCGGCGGTAGTGGTGGTGGTTCTGGG<br>GGTGGTAGCCATCACCACCATCACCAT | MKRVNAFNDLKRIGDDKVTAIGMGTWGIG<br>GRETPDYSRDKESIEAIRYGLELGMNLIDTA<br>EFYGAGHAEEIVGEAIKEFEREDIFIVSKVW<br>PTHFGYEEAKKAARASAKRLGTYIDLYLLH<br>WPVDDFKKIEETLHALEDLVDEGVIRYIGVS<br>NFNLELLQRSQEVMRKYEIVANQVKYSVKD<br>RWPETTGLLDYMKREGIALMAYTPLEKGTL<br>ARNECLAKIGEKYGKTAAQVALNYLIWEEN<br>VVAIPKASNKEHLKENFGAMGWRLSEEDR<br>EMARRCVGGGSGGGSGGGSHHHHHH |
| AdhD_MxTP | ATGAAGCGCGTTAACGCATTTAACGACCTG<br>AAACGTATTGGGGATGATAAGGTTACTGCA<br>ATTGGTATGGGGACATGGGGCATCGGGGG<br>CCGTGAAACTCCTGACTATTCCCGCGACAA<br>GGAGTCCATCGAAGCAATCCGGTATGGCTT<br>GGAGCTCGGCATGAACCTTATTGACACGGC<br>TGAATTTTATGGGGCGGGCCACGCAGAAG<br>AGATTGTTGGTGAGGCTATCAAGGAATTTG<br>AACGTGAGGACATTTTTATCGTGTCGAAGG<br>TATGGCCGACACATTTTGGGTACGAAGAAG<br>CAAAGAAAGCGGCCCGGGCCTCGGCCAAG<br>CGCTTGGGTACTTATATTGATTTGTACTTAT<br>TGCACTGGCCTGTTGACGATTTCAAAAAAA<br>TTGAAGAGACCTTGCATGCCCTTGAAGACC<br>TCGTCGACGAGGGGGTCATCCGTTACATC<br>GGCGTTTCAAACTTCAACTTAGAACTTCTCC<br>AACGGTCCCAGGAGGTGATGCGCAAATAC<br>GAGATCGTAGCGAACCAAGTGAAGTATTCT<br>GTAAAGGACCGGTGGCCAGAAACCACGGG<br>TCTCCTGGATTACATGAAGCGGGAGGGCAT<br>CGCTTTAATGGCTTACACGCCACTTGAGAA<br>AGGTACTTTGGCGCGGAATGAATGTCTTGC<br>GAAGATCGGGGAGAAATACGGTAAAACCG<br>CGGCACAGGTAGCCCTCAACTATCTGATTT<br>GGGAAGAAAATGTGGTGGCGATCCCTAAG<br>GCATCTAATAAGGAACACTTGAAAGAGAAT<br>TTCGGGGCCATGGGTTGGCGCCTTAGCGA<br>AGAAGACCGCGAGATGGCTCGTCGCTGTG<br>TTGGCGGCGGTAGTGGTGGTGGTTCTGGG | MKRVNAFNDLKRIGDDKVTAIGMGTWGIG<br>GRETPDYSRDKESIEAIRYGLELGMNLIDTA<br>EFYGAGHAEEIVGEAIKEFEREDIFIVSKVW<br>PTHFGYEEAKKAARASAKRLGTYIDLYLLH<br>WPVDDFKKIEETLHALEDLVDEGVIRYIGVS<br>NFNLELLQRSQEVMRKYEIVANQVKYSVKD<br>RWPETTGLLDYMKREGIALMAYTPLEKGTL<br>ARNECLAKIGEKYGKTAAQVALNYLIWEEN<br>VVAIPKASNKEHLKENFGAMGWRLSEEDR<br>EMARRCVGGGSGGGSGGGSPEKRLTVGS<br>LRR |

| | | |
|---|---|---|
| | GGTGGTAGCCCGGAGAAACGGCTCACTGT<br>AGGGAGCCTGCGTCGT | |
| PTDH_His | ATGTTGCCTAAGCTTGTGATTACACATCGC<br>GTACATGAGGAGATTCTGCAGTTATTGGCG<br>CCTCATTGTGAGCTGATTACCAACCAGACA<br>GATTCAACCCTCACGCGTGAAGAAATTTTA<br>CGCCGTTGTCGTGACGCTCAAGCGATGAT<br>GGCATTTATGCCGGACCGCGTTGATGCTGA<br>CTTCTTACAAGCCTGTCCTGAACTGCGTGT<br>AATCGGGTGCGCTCTTAAGGGGTTTGACAA<br>TTTCGATGTAGATGCCTGCACGGCTCGCGG<br>GGTATGGTTAACATTCGTCCCTGATCTCCT<br>GACGGTTCCAACCGCGGAGCTGGCAATTG<br>GCCTCGCGGTAGGTTTAGGGCGTCATCTG<br>CGGGCAGCAGACGCATTCGTCCGCTCGGG<br>CAAATTCCGTGGTTGGCAACCACGTTTCTA<br>TGGTACAGGGCTTGATAATGCTACGGTAGG<br>CTTCCTGGGTATGGGGGCCATCGGTCTTG<br>CCATGGCAGACCGTCTTCAGGGGTGGGGC<br>GCAACCTTACAGTATCATGAGGCTAAGGCT<br>TTGGATACGCAGACAGAGCAGCGTCTCGG<br>TCTGCGCCAAGTCGCGTGCAGCGAACTGTT<br>CGCTAGTTCGGATTTTATTCTGCTTGCGTTA<br>CCGCTCAATGCAGACACCTTGCATCTTGTA<br>AACGCTGAACTTCTGGCCTTGGTACGGCCT<br>GGCGCACTCCTCGTGAACCCTTGCCGCGG<br>GAGTGTTGTTGATGAGGCCGCGGTGCTTG<br>CTGCGCTTGAACGCGGCCAGCTGGGCGGC<br>TACGCTGCCGATGTGTTTGAAATGGAAGAC<br>TGGGCCCGCGCGGATCGTCCACAACAGAT<br>TGACCCGGCGCTGCTTGCACATCCAAATAC<br>GCTTTTCACTCCGCATATCGGCTCGGCGGT<br>ACGCGCCGTACGTCTTGAGATTGAGCGGT<br>GTGCTGCCCAGAACATTCTCCAAGCGCTGG<br>CCGGCGAGCGTCCTATCAATGCAGTCAATC<br>GCTTGCCGGGTGGTGGCAGCGGTGGGGG<br>CTCGGGTGGTGGTTCTCATCACCACCATCA<br>CCAT | MLPKLVITHRVHEEILQLLAPHCELITNQTDS<br>TLTREEILRRCRDAQAMMAFMPDRVDADF<br>LQACPELRVIGCALKGFDNFDVDACTARGV<br>WLTFVPDLLTVPTAELAIGLAVGLGRHLRAA<br>DAFVRSGKFRGWQPRFYGTGLDNATVGFL<br>GMGAIGLAMADRLQGWGATLQYHEAKALD<br>TQTEQRLGLRQVACSELFASSDFILLALPLN<br>ADTLHLVNAELLALVRPGALLVNPCRGSVV<br>DEAAVLAALERGQLGGYAADVFEMEDWAR<br>ADRPQQIDPALLAHPNTLFTPHIGSAVRAV<br>RLEIERCAAQNILQALAGERPINAVNRLPGG<br>GSGGGSGGGSHHHHHH |
| PTDH_MxTP | ATGTTGCCTAAGCTTGTGATTACACATCGC<br>GTACATGAGGAGATTCTGCAGTTATTGGCG<br>CCTCATTGTGAGCTGATTACCAACCAGACA<br>GATTCAACCCTCACGCGTGAAGAAATTTTA<br>CGCCGTTGTCGTGACGCTCAAGCGATGAT<br>GGCATTTATGCCGGACCGCGTTGATGCTGA<br>CTTCTTACAAGCCTGTCCTGAACTGCGTGT<br>AATCGGGTGCGCTCTTAAGGGGTTTGACAA<br>TTTCGATGTAGATGCCTGCACGGCTCGCGG<br>GGTATGGTTAACATTCGTCCCTGATCTCCT<br>GACGGTTCCAACCGCGGAGCTGGCAATTG<br>GCCTCGCGGTAGGTTTAGGGCGTCATCTG<br>CGGGCAGCAGACGCATTCGTCCGCTCGGG<br>CAAATTCCGTGGTTGGCAACCACGTTTCTA<br>TGGTACAGGGCTTGATAATGCTACGGTAGG<br>CTTCCTGGGTATGGGGGCCATCGGTCTTG<br>CCATGGCAGACCGTCTTCAGGGGTGGGGC<br>GCAACCTTACAGTATCATGAGGCTAAGGCT<br>TTGGATACGCAGACAGAGCAGCGTCTCGG<br>TCTGCGCCAAGTCGCGTGCAGCGAACTGTT<br>CGCTAGTTCGGATTTTATTCTGCTTGCGTTA<br>CCGCTCAATGCAGACACCTTGCATCTTGTA<br>AACGCTGAACTTCTGGCCTTGGTACGGCCT<br>GGCGCACTCCTCGTGAACCCTTGCCGCGG<br>GAGTGTTGTTGATGAGGCCGCGGTGCTTG<br>CTGCGCTTGAACGCGGCCAGCTGGGCGGC<br>TACGCTGCCGATGTGTTTGAAATGGAAGAC<br>TGGGCCCGCGCGGATCGTCCACAACAGAT<br>TGACCCGGCGCTGCTTGCACATCCAAATAC | MLPKLVITHRVHEEILQLLAPHCELITNQTDS<br>TLTREEILRRCRDAQAMMAFMPDRVDADF<br>LQACPELRVIGCALKGFDNFDVDACTARGV<br>WLTFVPDLLTVPTAELAIGLAVGLGRHLRAA<br>DAFVRSGKFRGWQPRFYGTGLDNATVGFL<br>GMGAIGLAMADRLQGWGATLQYHEAKALD<br>TQTEQRLGLRQVACSELFASSDFILLALPLN<br>ADTLHLVNAELLALVRPGALLVNPCRGSVV<br>DEAAVLAALERGQLGGYAADVFEMEDWAR<br>ADRPQQIDPALLAHPNTLFTPHIGSAVRAV<br>RLEIERCAAQNILQALAGERPINAVNRLPGG<br>GSGGGSGGGSPEKRLTVGSLRR |

| | |
|---|---|
| | GCTTTTCACTCCGCATATCGGCTCGGCGGT ACGCGCCGTACGTCTTGAGATTGAGCGGT GTGCTGCCCAGAACATTCTCCAAGCGCTGG CCGGCGAGCGTCCTATCAATGCAGTCAATC GCTTGCCGGGTGGTGGCAGCGGTGGGGG CTCGGGTGGTGGTTCTCCAGAAAAGCGGC TTACTGTGGGGTCGTTACGGCGT |

**Supplementary Table 2. DNA primers used in this study.**

| Primer | Sequence (5’->3’) |
|---|---|
| pETDuet_F | ATTAACCTAGGCTGCTGCCACCG |
| pETDuet_B1 | TGTATATCTCCTTCTTATACTTAACTAATATACTAAGATGGGGAATTG |
| pETDuet_B2 | GTATATCTCCTTCTTAAAGTTAAACAAAATTATTTCTAGAGGGG |
| Mx_pmut_F | CTTATATTCCCAGTTACATCGGGGTGGCGAGATCGAGACAATTCGCC |
| Mx_pmut_R | CCGATGTAACTGGGAATATAAGC |
| NanoLuc_His_F | AAGTATAAGAAGGAGATATACAATGGTTTTCACCTTAGAAGATTTCGTG |
| NanoLuc_His_R | GCAGCAGCCTAGGTTAATTCAATGGTGATGGTGGTGATGTGAACCGCCCCCGCTC |
| Adhd_His_F | AAGTATAAGAAGGAGATATACAATGAAGCGCGTTAAC |
| Adhd_His_R | GCAGCAGCCTAGGTTAATTCAATGGTGATGGTGGTGATGGCTACCACCCCCAGAAC |
| PTDH_His_F | TTAACTTTAAGAAGGAGATATACATGTTGCC |
| PTDH_His_R | GCAGCAGCCTAGGTTAATTCAATGGTGATGGTGGTGATGAGAACCACCACCCGAGC |

**Supplementary Table 3. Cryo-EM data collection, refinement, and validation statistics.**

| | Mx_pmut T=1 (EMDB-44383) (PDB 9B9I) | Mx_pmut T=3 (EMDB-44388) (PDB 9B9Q) | Mx_pmut T=4 (EMDB-44427) (PDB 9BC8) |
|---|---|---|---|
| **Data collection and processing** | | | |
| Magnification | 45,000x | 45,000x | 45,000x |
| Voltage (kV) | 200 | 200 | 200 |
| Electron exposure (e–/Å$^2$) | 39.18 | 39.56 | 39.56 |
| Defocus range (μm) | -1.0 to -1.8 | -1.0 to -1.8 | -1.0 to -1.8 |
| Pixel size (Å) | 0.91 | 0.91 | 0.91 |
| Symmetry imposed | I | I | I |
| Initial particle images (no.) | 66,813 | 37,600 | 37,600 |
| Final particle images (no.) | 61,808 | 12,967 | 4,386 |
| Map resolution (Å) | 2.86 | 3.14 | 3.46 |
| FSC threshold | 0.143 | 0.143 | 0.143 |
| **Refinement** | | | |
| Initial model used (PDB code) | | | |
| Model resolution (Å) | 3.2 | 3.6 | 3.9 |
| FSC threshold | 0.5 | 0.5 | 0.5 |
| Map sharpening *B* factor (Å$^2$) | -126.4 | -93.1 | -87.9 |
| Model composition | | | |
| Non-hydrogen atoms | 2,038 | 6,569 | 8,740 |
| Protein residues | 262 | 848 | 1,128 |
| Ligands | 0 | 0 | 0 |
| *B* factors (Å$^2$) | | | |
| Protein | 43.61 | 60.14 | 104.39 |
| R.m.s. deviations | | | |
| Bond lengths (Å) | 0.005 | 0.005 | 0.005 |
| Bond angles (°) | 1.093 | 1.039 | 1.095 |
| Validation | | | |
| MolProbity score | 2.09 | 2.30 | 2.04 |
| Clashscore | 8.64 | 11.77 | 13.43 |
| Poor rotamers (%) | 3.33 | 2.94 | 3.41 |
| Ramachandran plot | | | |
| Favored (%) | 96.51 | 94.86 | 93.97 |
| Allowed (%) | 3.49 | 5.14 | 5.94 |
| Disallowed (%) | 0 | 0 | 0.9 |
